# Single-cell chromatin and transcriptome dynamics of Synovial Fibroblasts transitioning from homeostasis to pathology in modelled TNF-driven arthritis

**DOI:** 10.1101/2021.08.27.457747

**Authors:** Marietta Armaka, Dimitris Konstantopoulos, Christos Tzaferis, Matthieu D Lavigne, Maria Sakkou, Anastasios Liakos, Petros P Sfikakis, Meletios A Dimopoulos, Maria Fousteri, George Kollias

**Affiliations:** Institute for Fundamental Biomedical Research, Biomedical Sciences Research Center “Alexander Fleming”, Vari, Greece; Institute for Bioinnovation, Biomedical Sciences Research Center “Alexander Fleming”, Vari, Greece; First Department of Propaedeutic Internal Medicine, National and Kapodistrian University of Athens Medical School, Athens, Greece; Center of New Biotechnologies & Precision Medicine, National and Kapodistrian University of Athens Medical School, Athens, Greece; Joint Rheumatology Program, National and Kapodistrian University of Athens Medical School, Athens, Greece; Department of Clinical Therapeutics, National and Kapodistrian University of Athens Medical School, Athens, Greece; Department of Physiology, Medical School, National and Kapodistrian University of Athens, Athens, Greece

## Abstract

Synovial fibroblasts (SFs) are specialized cells of the synovium that provide nutrients and lubricants for the maintenance of proper function of diarthrodial joints. Chronic TNF signals are known to trigger activation of SFs and orchestration of arthritic pathology via proinflammatory effector functions, secretion of cartilage degrading proteases and promotion of osteolysis. We performed single-cell (sc) profiling of SF’s transcriptome by RNA-sequencing (scRNA-seq) and of chromatin accessibility by scATAC-seq in normal mouse SFs and SFs derived from early and advanced TNF-driven arthritic disease. We describe here distinct subsets of SFs in the homeostatic synovium, serving diverse functions such as chondro- and osteogenesis, tissue repair and immune regulation. Strikingly, development of spontaneous arthritis by transgenic TNF overexpression primes the emergence of distinct pathology-associated SF subtypes. We reveal 7 constitutive and 2 disease-specific SF subtypes. The latter emerge in the early stage, expand in late disease and are localized in areas at the interface between the invasive pannus and the articular bone. The associated transcription profiles are characterized by enhanced inflammatory responses, promigratory behaviour, neovascularization and collagen metabolic processes. Temporal reconstruction of transcriptomic events indicated which specific sublining cells may function as progenitors at the root of trajectories leading to intermediate subpopulations and culminating to a destructive lining inflammatory identity. Integrated analysis of chromatin accessibility and transcription changes revealed key transcription factors such as Bach and Runx1 to drive arthritogenesis. Parallel analysis of human arthritic SF data showed highly conserved core regulatory and transcriptional programs between the two species. Therefore, our study dissects the dynamic SF landscape in TNF-mediated arthritis and sets the stage for future investigations that might address the functions of specific SF subpopulations to understand joint pathophysiology and combat chronic inflammatory and destructive arthritic diseases.

## Introduction

Chronic arthritides including Rheumatoid Arthritis (RA) are complex inflammatory disorders that primarily affect diarthrodial joints causing high morbidity and mortality in human patients. Cells driving pathogenicity in the affected joints, include an expanding mass of synovial fibroblasts (SFs) typically infiltrated by myeloid and lymphoid cells, which together contribute to the development of an invasive pannus that degrades cartilage and promotes osteolysis^1,2^. Early studies in transgenic mice have established a key role for TNF in driving the full pathogenic process ^3,4^. This was confirmed later in humans by the introduction of anti-TNF therapies that proved efficacious in neutralizing disease in a large percentage of rheumatoid arthritis (RA) patients ^5^. Further genetic studies in murine arthritis models revealed that, in particular, TNF signaling in synovial fibroblasts (SFs) mediates persistent fibroblast activation and promotes pro-proliferative, immune-regulatory and invasive characteristics. These functions are both necessary and sufficient to orchestrate initiation and progression of the inflammatory and damaging pathology even in the absence of adaptive immune responses^6–8^ qualifying SFs as key effector cells and crucial therapeutic target in chronic arthritis.

SFs are the major cellular component of the synovial membrane, a highly specialized, multifunctional connective tissue membrane comprised of two anatomically distinct layers: lining SFs (LSFs) and the recently identified CXCR3+ lining macrophages^9^ that form a thin outer layer adjacent to inmost structures consisting of sublining SFs (SLSFs), macrophages, adipose cells, nerves and blood vessels^10^. SFs in the RA proinflammatory microenvironment acquire an aggressive phenotype, reminiscent of transformed migratory tumor-like cells^11^. They operate as immune-modulatory cells by secreting cytokines and chemokines and mediate cartilage destruction by over-expressing MMP1, MMP3 and MMP9 matrix metallo-proteases^12,13^ as well as the receptor activator of NF-κB ligand (RANKL/*Tnfsf11*), which causes excessive osteoclastogenesis leading to bone erosions^14,15^.

Histopathological analysis of RA joints, and studies using a combination of single-cell (sc) and bulk RNA-seq analysis of RA patients, indicated that distinct fibroblasts subpopulations in the lining and the sublining synovial compartments are linked to specific disease features. Lining fibroblasts markers include, podoplanin (PDPN) and Lubricin/Proteoglycan 4 (PRG4), whereas sublining SFs are characterized by high THY1 and PDPN expression. The RA SF subpopulations are characterized by differential expression of several markers such as CD34, VCAM1, FAP and proinflammatory mediators, such as CXCL12, CCL2 and IL6^16,17^. More recent studies classified the fibroblasts found in the synovial lining zone as being predominantly responsible for driving articular damage whereas fibroblasts located in the sublining layer express genes that function towards inflammation^18,19^. Additional recent evidence revealed a dominant Notch-mediated interplay of perivascular SLSFs with endothelial cells, establishing a positional gradient of Thy1^hi^ SLSFs towards Thy1^low^/Prg4^hi^ LSFs and driving tissue inflammation^20^. Although these studies were instrumental in providing valuable insights in the classification of pathogenic SF subpopulations and their associated functions in the RA synovium, the homeostatic to pathological transitions of SFs and the molecular networks that drive them have remained unclear.

In this study, we aimed to uncouple and characterize the homeostatic and pathological functions of SFs in the human TNF overexpressing (*hTNFtg*) mouse model^3^. We undertook an integrative approach by combining sc transcriptomic and chromatin accessibility data to define the underlying molecular switches that determine the staging and progression of disease from healthy to early inflammatory and subsequent destructive synovial tissue. Our data reveal the early emergence and further expansion of distinct pathogenic SF subtypes characterized by specific differentially activated pathways and regulatory networks emanating from a progenitor state that appear repressed in the normal sublining synovium. Changes observed in SFs transcriptome were highly correlating to chromatin accessibility alterations and cellular trajectory inference pinpointed to novel key transcription factors and target genes driving the expansion of the pathogenic clusters at specific time and locations during disease progression. Lastly, integrative meta-analysis of our murine data with available human RA data revealed a highly conserved core regulatory transcriptional program, validating our modelling approach and revealing a set of novel biomarkers specific to TNF-driven RA. Our results provide a solid translational potential to prioritize novel molecular and cellular targets specific for the pathogenic transitions of synovial fibroblasts in RA.

## Results

### Multi sc-omic analysis of *hTNFtg* mouse model of chronic inflammatory polyarthritis

To characterize disease progression and pinpoint what differentiates homeostasis from pathogenesis at the level of SF subpopulations in joints synovium, we integrated sc transcriptomic and chromatin accessibility profiles (Figure 1a). Our setup included cells from healthy tissue (WT, 4 weeks of age (n=3)), *hTNFtg* mice at an early disease stage displaying synovial inflammation (*hTNFtg*/4, 4 weeks of age (n=3)), and at an established pathological stage displaying pannus formation, inflammation, cartilage and bone damage) (*hTNFtg*/8, 8 weeks of age (n=3) (Suppl. Figure 1a). 3’ single cell mRNA-sequencing libraries (10X Genomics) were generated for 6,667 sorted non-hematopoietic stromal cells (Cd45^-^, Cd31^-^, Ter119^-^, Pdpn^+^) isolated from whole ankle joint synovium (Figure 1a and Suppl. Figure 1b). In parallel, single-cell assay for transposase-accessible chromatin using sequencing (scATAC-seq, 10X Genomics) was performed to determine the chromatin accessibility landscape across 6,679 single nuclei (Pdpn-, Thy1+ Fibroblasts have been excluded during gating and sorting (Suppl. Figure 1b). Healthy and *hTNFtg* cells were pooled in each experimental modality, to create a common baseline between homeostatic and pathogenic conditions.

**Figure 1.**
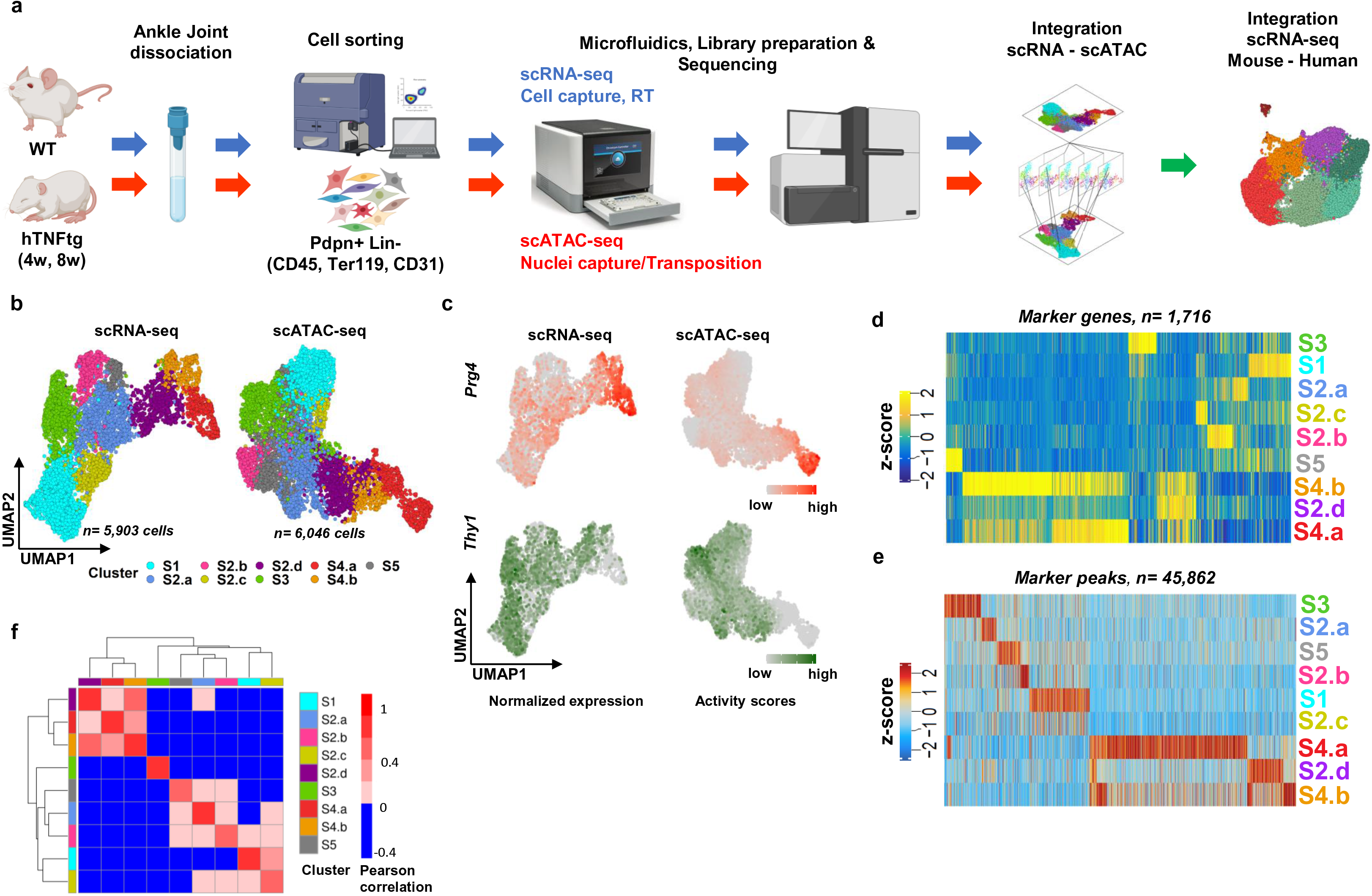
Multiomic transcriptional and epigenetic single cell analysis of Fibroblastic Synovial cells. (a). Schematic representation of the experimental workflow. We collected ankle synovial tissue from WT and hTNFtg mice, enzymatically disaggregated the tissue and sorted the cells into one gate representing fibroblasts (CD45-, Ter119-, CD31-, Pdpn+). We profiled the cells with both sc 3’ mRNAseq (scRNA-seq) and scATACseq using 10X technology and performed cross-modalities (scRNA-seq-scATAC-seq) as well as cross-species scRNA-seq integrative analyses with publicly available human RA datasets. (b). High quality filtered Synovial Fibroblasts (SFs) (n=5,903 cells for the sc-RNAseq and n=6,046 nuclei for the sc-ATACseq) projected in UMAP space and colored by cluster. (c). Feature plots of the SFs shown in panel b, displaying normalized expression values (for scRNA-seq) and activity scores (for scATAC-seq) for Prg4 and Thy1 genes. (d). Heatmap showing the unsupervised clustering of scRNAseq marker genes (up-regulated genes in at least one subpopulation vs the others) according to their average scaled expression values (z-score). (e). Heatmap showing the unsupervised clustering of scATACseq marker peaks (regions with up-regulated accessibility in at least one subpopulation vs the others) according to their average scaled activity scores (z-score) (see Methods). (f). Heatmap of Pearson correlation coefficients between average scaled expression values(RNA) and average scaled activity scores(ATAC) of the most variable genes identified by scRNA-seq analysis.

Inspection of marker genes of the major clusters that were isolated by FACS, was used to annotate the respective cell types (Methods) and suggested that 10% of the sequenced cells/nuclei represented non-fibroblastic cells such as: osteoblasts (*Alpl*, *Bglap2*, *Ostn*), chondrocytes (*Chad*, *Clip2*), myoblasts/myocytes (*Des, Actn3*) and vascular cells (*Cdh5*) (Suppl. Figure 2a, b). ScATAC-seq cluster annotation was performed using canonical correlation analysis (CCA) and enabled to match scRNA-seq and scATAC-seq cluster identities (Suppl. Figure 3a, b and Methods).

We focused on the 5,903 and 6,046 cells/nuclei presenting SFs characteristics in scRNA-seq and scATAC-seq respectively (Suppl. Figures 2c, 3c). Sub-clustering analysis of SF-specific molecular maps resolved nine fibroblastic clusters: S1, S2 (a, b, c and d), S3, S4 (a and b), and S5 (Figure 1b and Suppl. Figures 2d, 3d). Using as a proxy the classical markers Prg4 and Thy1^16,18,19^, we observed a compartmentalization of LSLs (S4a, (Prg4+)) vs SLSFs (S1, S2a, b, c, S3, S5 (Thy1+)) (Figure 1c). We also noted that clusters S2d and S4b, which are mainly present in disease stage (*hTNFtg*/4,8), expressed both genes (Suppl. Figure 4). Clustering of cells based on all 1,716 scRNA-seq marker genes expression (*i.e.* up-regulated in at least one cluster vs the others) and on accessibility patterns at 45,862 marker peaks (i.e. with increased accessibility in at least one SF subpopulation compared to the others) revealed cell specificity and shared patterns both at transcriptional and chromatin levels (Figure 1d, e). In fact, high correlation coefficient scores between gene expression and chromatin accessibility were observed not only within clusters, but also across clusters and suggested some architectural/functional overlap amongst SLSFs and amongst LSFs and S2d, S4b clusters (Figure 1f). Overall, by using a combined-omics approach we could deconvolve SFs varieties. Hence, we reveal specific patterns of gene expression and the associated chromatin accessibility signatures, which may be used to further characterize RA molecular markers and to understand the gene regulatory networks driving its pathophysiology (see below).

### High-resolution maps of transcription regulation in homeostatic joints

We first characterized SF populations RNA expression specificities in healthy homeostatic joints by looking at WT SFs independently (Figure 2a) and by quantifying the number of cells detected per sample and per cluster (Figure 2b). We detected cluster specific up-regulated genes (Suppl. Figure 5a), which corresponded to established SFs marker genes, and we also identified genes that so far had not been linked to SFs biology, probably given their limited expression in a few specific cells (Suppl. Table 1).

**Figure 2.**
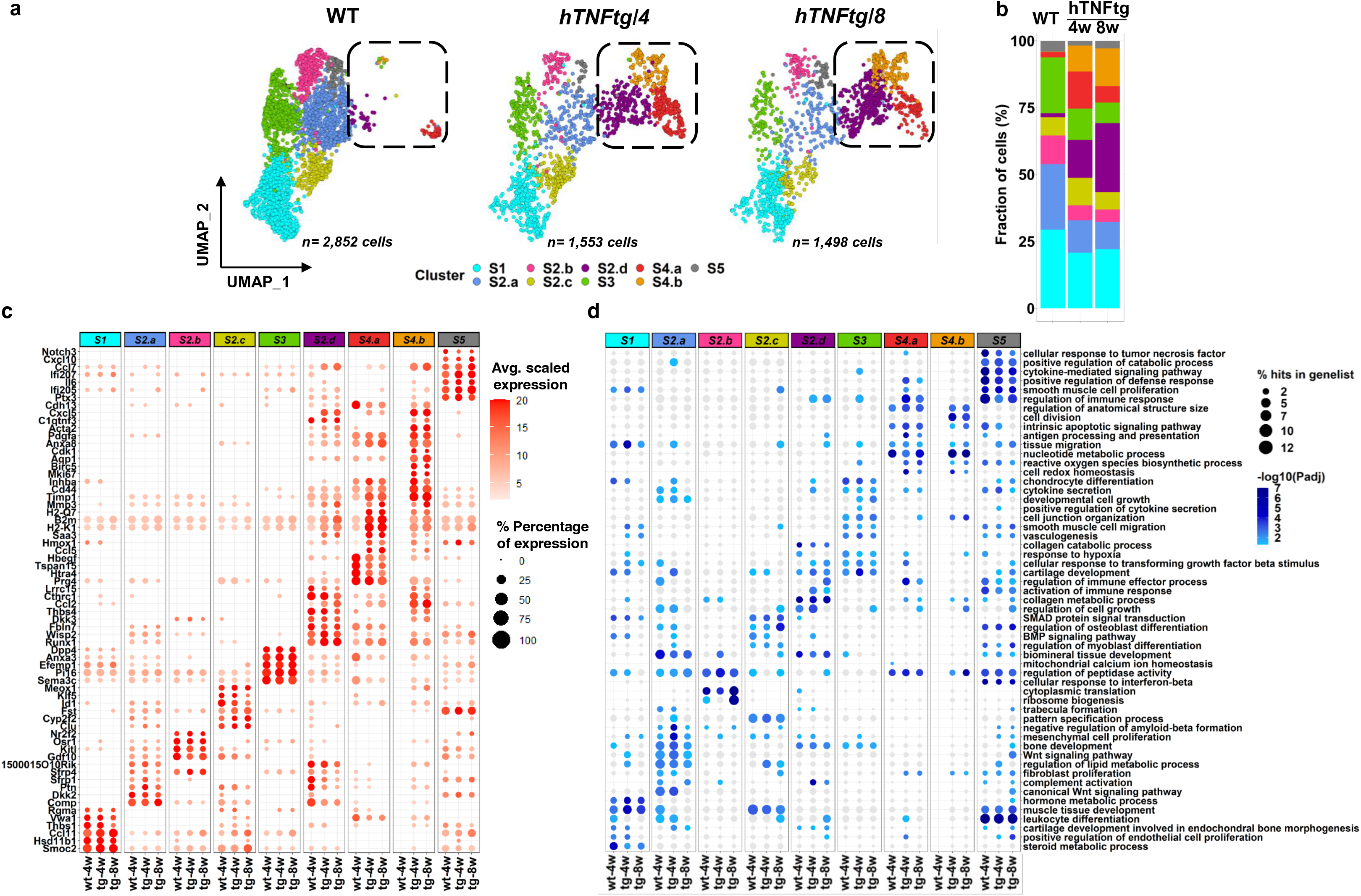
Functional remodeling of the synovial mesenchyme between WT healthy and hTNFtg arthritic joints. (a). UMAP representation of scRNA-seq-derived SFs distributions for the three samples separately (WT, hTNFtg/4 weeks and hTNFtg/ 8 weeks as indicated). The cells are colored by cluster identities and the marked area highlights the dynamic changes of the intermediate (I) and lining (L) subpopulations during disease progression. (b). Stacked barchart showing relative abundances (%) of clusters across samples (WT, hTNFtg/4 weeks and hTNFtg/ 8). (c). Dotplot of a selection of cluster marker genes identified by scRNA-seq. The color of the dot shows the intensity of expression while the size denotes the percentage of cells expressing the gene in each cluster and condition. (d). Dotplot of a selection of representative gene ontologies (GO, Biological Processes) per clusters/samples as identified by Functional enrichment analysis (see Methods). The colour of the dot shows the statistical significance (-Log10(Padj)) while the size denotes the percentage of the cluster marker genes found in the GO gene list.

S4a expressed high levels of *Prg4* (Prg4^high^) without Thy1, as well as other genes previously reported as markers of the lining phenotype, such as *Tspan15*, *Hbegf* and *Htra4* ^18,19^ (Figure 2c). In WT joints, these LSFs were clearly demarcated from the sublining cells (Thy1+, Prg4-) because of the very limited number of S2d and S4b cells (Thy1+ Prg4+) (Figure 2a, b), a result that establishes that compartmentalization of Thy 1 and Prg4 expression is more robust in WT tissues. Functional enrichment analysis revealed that, in contrast to the reported destructive profile of the lining cluster in arthritic disease^18^, in normal conditions LSFs tend to preserve tissue homeostasis by regulating the responses towards oxidative stress and the related induced cell death, as well as the homeostasis of mitochondrial calcium, a fundamental signaling modulator^21^ (Figure 2d).

Regarding Thy1^+^ SLSF populations, we find that S1 transcriptional state is marked by the expression of *Smoc2, Thbs1, Vwa1* genes that encode matricellular proteins and the BMP co-receptor *Rgma* (Figure 2c and Suppl. Table 1, 2), which indicate a strong chondrogenic potential confirmed by the activated BMP/SMAD signaling pathways detected in the GO enrichment analysis (Figure 2e). The expression of genes associated with steroid metabolism, including the cortisone-conversion enzyme *Hsd11d1* provides to S1 cells a potential anti-inflammatory role^22^.

S2 SF subtypes are characterized by common (*Comp*, *Ptn*, *Gdf10)* and divergent marker genes and functions (Fig 2c, d and Suppl Table 1, 2). In particular, the S2a population is defined by the high expression of WNT modulators *Dkk2* and *Sfrp1,* in accord to the GO enrichment in WNT-mediated responses, TGF activity and osteogenesis. In addition, the specific expression of Ecrg4 gene indicates a role of S2a in regulating tissue repair processes (wound healing)^23^. In S2b, gene expression is correlated with joint morphogenesis and reparative processes; *e.g. Osr1* regulates *Prg4*^24^ and plays a pivotal role in fibroblast differentiation^25^. Moreover, *Nr2f2* (COUP-TFII) marker gene is implicated in cell fate decisions of stem cells^26^. Along the same line of evidence, S2c cellular state is characterized by BMP signaling pathway activation, suggesting S2c involvement in synovium renewal and protection. The role of S2c in tissue homeostasis is also supported by specific expression of *Klf5* and *Clu*, involved in regulating proliferation, senescence and/or responses to oxidative stress^27–29^. Other signaling modulators highly expressed in S2c, such as *Id1* and *Meox1,* are associated with increased expression of the p16/Ink4a locus^30–32^, a cell cycle regulator previously suggested as a therapeutic target for RA^33^.

The gene expression signature of S3 suggests that these cells drive processes relative to vasculogenesis and regulation of type 2 immune responses, as characterized by the expression of Pi16, which functions in pain and fibroblast/endothelial crosstalk^34^, of the physiological vascular normalizing modulators *Sema3c*^35^ and *Efemp1*^36,37^ as well as of the glucose and immune regulator *Dpp4*^38^ (Figure 2c, d and Suppl. Table 1, 2).

Finally, S5 cells show activation of cytokines and chemokine pathways (*Ccl7*, *Cxcl10, IL6* and *Ptx3*) and are associated with immune-regulatory functions including response to IFN-beta/gamma and LIF, indicating a strong immunoregulatory role in the synovial membrane under healthy conditions. Notably, Notch3, a gene recently highlighted for its role in driving SF identity in the perivascular/sublining layer of arthritic synovium^20^, is also expressed in normal conditions exclusively in cluster S5 (Figure 2c and Suppl. Table 1, 2).

Overall, the analysis of SFs in naïve conditions highlights a previously underexplored functional diversity underlying the homeostasis of synovial membrane.

### Development of inflammatory arthritis associates with transcriptional remodeling of SF populations and functions

We next sought to dissect the processes underlying the appearance and maintenance of TNF-induced pathological states of SFs. Differential abundance analysis revealed disease-enriched cell subpopulations: S2d and S4b. Notably, the proportion of S2d cells increased from almost undetectable levels (2%) in healthy conditions to 25% in the *hTNFtg* joints (Figure 2a, b). Similarly, S4b class, which was virtually absent in WT (0.17%), became gradually more evident during arthritogenesis (9.72% and 14.08% in early and established arthritis respectively) (Figure 2a, b). The mixed expression signature of Prg4 and Thy1 (Prg4+Thy1+), which characterize this “intermediate” group of cells, is thus a strong marker of disease state that is observed mainly in *hTNFtg* conditions (Figure 1c, 2a, Suppl. Figure 4).

In fact, correlation analysis on the most variable genes (MVGs) (Suppl. Figure 5b) of SFs clusters highlighted a striking overlap in the transcriptional profiles of the Prg4^High^ S4a SF and the intermediate S4b and S2d SF subpopulations, which was already suggested from the patterns of selected representative marker genes and GOs (see Figure 2c, d). Correlation scores were higher between *hTNFtg* cells indicating an acute and stable change in these SFs expression signatures after onset of arthritis (Suppl. Figure 5b). Intra-cluster differential expression analysis (DEA) between *hTNFtg* and WT cells identified what changes occurred after disease onset and how they were mainly detected in S2d, S4a and S4b SFs (Suppl. Figure 5c). The significant intersection of cluster and disease marker genes (inter-intra DE) for intermediate and lining clusters suggests their synergism and points to what genes are probably essential to drive pathogenic functions (Suppl. Figure 5c-e, Suppl. Table 2 and see below). The gain in intermediate cells was offset by the shrinkage in proportion of the number of other cell types S2a, b, c, S3 and, to a lesser degree, S1 and S5 (Figure 2a, b) and these clusters showed a more homogenous signature and less DE genes between WT and *hTNFtg*, thus underlying their common and stable functions in healthy and disease joints (Suppl. Figure 5b, c).

Analysis of the shared disease specific up-regulated genes (Suppl. Figure 5a-right, c and d) suggest how they might drive the extensive rearrangement of synovium functions and properties during arthritis (Suppl. Figure 5e, and see below). For instance, S2d cells express highly important genes for joint pathology (Figure 2c, d) including: the ECM component *Fbln7*^39^, which was recently shown to regulate calcium deposition in diseased kidney tissues, suggesting its role in arthritic bone remodeling ^40^; the matricellular protein *Thbs4*; the vascular remodeler *Cthrc1*, which has also been proposed as a marker for embryonic progenitors of SFs, fibrocartilage cells of the enthesis^41^ and fibrotic lung fibroblasts ^42^, and is currently tested as a diagnostic marker for RA (https://clinicaltrials.gov/ct2/show/NCT04092244). S2d SFs also express *Lrrc15*, a recently identified marker for cancer associated fibroblasts (CAFs) and activated fibroblasts^43,44^. Finally, expression of *Dkk3*, a miscellaneous member of DKK family that possibly regulates TGFbeta rather than WNT pathway^45^, associates the murine S2d transcriptional state with the previously described human SC-F3 (DKK3+) SF cluster^19^. In accord, we find that the biological processes characterising S2d SFs extend from the regulation of immune and redox response to cell fate determination and ECM remodeling, indicating a multi-potent transcriptional signature. In this respect, we highlight Runx1 transcription factor (TF) as an outstanding potential regulatory gene highly expressed in S2d cells. Although Runx1 has been mainly implicated in hematopoiesis and in the commitment/differentiation from chondroprogenitor cells into the chondrogenic lineage^46^, it could also function as a regulator of stromal cells transition from fibroblasts to activated myofibroblasts^47^.

S4b marker genes including *Mki67*, *Pdgfa*, *Birc5*, *Aqp1* suggest pathogenic functions associated with the processes of proliferation, cell cycle regulation and apoptosis (Figure 2c, d and Suppl. Figure 6a). Expression of cytoskeletal-related genes such as Acta2 also indicates that S4b cells display properties similar to those of a myofibroblast-like state of activated SFs. Finally, finding *C1qtnf3* adipokine, and other chemokines such as *Cxcl5*, as well as adhesion molecules such as *Cdh13* reinforces the idea that these cells probably largely contribute to the inflammatory process in arthritis (Figure 2c, d).

Analysis of the S4a lining SFs revealed that during TNF-mediated arthritis they preserve some of their homeostatic LSFs marker gene identity, but also show an expansion in the diversity of their transcriptome indicating that their reparative functions might be affected after disease onset. Indeed, we detected markers of inflammatory response (*Ccl2, Ccl5, Hmox1 Saa3*), Class I antigen presentation (*H2-K1, B2m, H2-Q7*) and ECM remodeling (*Mmp3, Timp1, Cd44*) (Figure 2c, d), in agreement with previous reports on arthritic LSFs^18,19^. Notably, a meticulous sub-clustering analysis of the S4a cluster indicated the presence of two groups of cells (subclusters hS4a (homeostatic) and iS4a (inflammatory)), where homeostatic gene expression and functions are minimized during disease expansion, while in parallel there is emergence and expansion of an aggressive inflammatory state (Suppl. Figure 7 and Suppl. Table 3).

To determine if the functional specialization of SFs described above is reflected in joints’ tissue architecture, we spatially localized the expression of some representative cluster specific marker genes in healthy and arthritis joints by immunofluorescence (IF) and confocal microscopy. S2a/b-associated marker Gdf10 and the S2a marker Comp were detected in Thy1-expressing mesenchymal cells, directly adjacent to the lining outermost cellular layer closer to the cartilage (Suppl. Figure 8a). Comp was also detected on chondrocytes on the articular cartilage. The S5 marker Notch3 was restricted to a smaller Thy1^+^ SLSF subpopulation in close proximity to the inner compartment of the synovial membrane (Suppl. Figure 8a). Furthermore, CD44 and Ki67, markers of the disease-expanding ‘intermediate’ clusters S4b and S2d, display a partial co-localization with Thy1 expressing SL fibroblasts (Suppl. Figure 8b/e). In particular, a specific spatial trend was identified for S2d and S4b SFs *as* Wisp2, Prg4, Mki67 and Cd44 markers localized in areas at the interface between the pannus and the articular bone and in the pannus tissue that has invaded the cartilage and bone during inflammation. In contrast S5 and S2 clusters are excluded from areas of bone erosion and restricted in the pannus tissue surrounding the joint colocalizing with Thy1 expression (Suppl. Figure 8a). Finally, Prg4 expressing SFs localization validated the S4a lining subpopulation directly adjacent to the articular cartilage and bone, and we observed in situ the expansion and invasive function of intermediate cells at the established disease state (Suppl. Figure 8c).

Collectively, these findings establish detailed molecular, functional, and anatomical maps charting the dynamic and diverse effects of TNF on the development and progression of the pathogenic SF states.

### Predictive markers of the inflammatory expansion of SFs in TNF-mediated arthritis

To test if our scRNA-seq results can be used to predict reliable arthritis marker gene expression in tissues, we performed bulk RNA-seq on sorted LSFs and SLSFs. LSF (CD31Cd31-, CD45-, Ter119-, CD90-, Pdpn+) and SLSF (CD31-, CD45-, Ter119-, CD90+, Pdpn+) populations of WT, *hTNFtg* 4 and 8 week-old mice were segregated and showed a clear separation of WT and *hTNFtg* cells (Suppl. Figure 9a). DEA revealed more changes in gene expression between lining and sublining SFs in healthy animals compared to *hTNFtg* counterparts (Suppl. Figure 9b, top, and Suppl. Table 4), again suggesting that SFs tend to lose their sharp bi-modal (lining vs sublining) character in arthritic tissues. By calculating FCs between LSF and SLSF for marker genes identified with scRNA-seq, we confirmed how S4a genes fit with a lining state signature, while S2d and S4b genes are more equally expressed in both states and the remaining clusters tend to be defined by genes up-regulated in SLSF state (Suppl. Figure 9d). Corroboratively, we find that more genes of sc clusters S4a are detected as bulk LSF markers, more genes representative of S5, S2b and S2b are bulk markers of SLSFs, while a balanced number of S2d and S4b markers are found in LSF and SLFs bulk DEG lists (Suppl. Figure 10a) and we highlight some representative genes for the SL, intermediate (I), and L cells (Suppl. Figure 10b). Finally, we found candidate diagnostic genes that loose or gain differences between LSF and SLSF during disease and propose that they can be used to test disease severity by sorting of SFs from joints biopsies followed by quantification of gene expression by qPCR (Suppl. Figure 10c).

### Development of arthritic pathology depends on activated epigenomic states in SFs

To identify the pathogenic molecular master switches that remain repressed in healthy joints and are activated in arthritis we also analyzed scATAC-seq data to find condition- and cell-type specific chromatin signatures and explore what TF and target genes are controlled at the epigenomic level. Accessible chromatin patterns recapitulated the significant expansion of the SFs subpopulations S2d and S4b upon disease progression (Figure 3a, b). Moreover, by performing a two-level differential accessibility (DA) analysis we characterized how SFs functional outputs are controlled by the variable repertoire of open chromatin regions (OCRs). We established first what genomic regions gained or lost scATAC-seq reads between given SF subtypes (inter-cluster analysis) (Figure 3c) and then which loci changed their status (opening or closing) in a given cluster in arthritic (*hTNFtg* 4weeks and *hTNFtg* 8 weeks combined) vs healthy (WT) SFs (intra-cluster analysis) (Figure 3d). We noted a particularly elevated number of cell- and subtype-specific OCRs in intermediate cells S2d, S4b and lining S4a and a striking gain in DNA accessibility at 27,9 and 49,8% at S2d and S4b specific loci (Figure 3d). Our data supports the idea that a drastic rearrangement of chromatin underlies the expansion of these cell types and we suggest that the identified regions can serve as novel disease-specific accessibility signatures (Figure 3c-d) and they could provide new mechanistic links to what/how key regulatory genes control arthrogenesis (see below).

**Figure 3.**
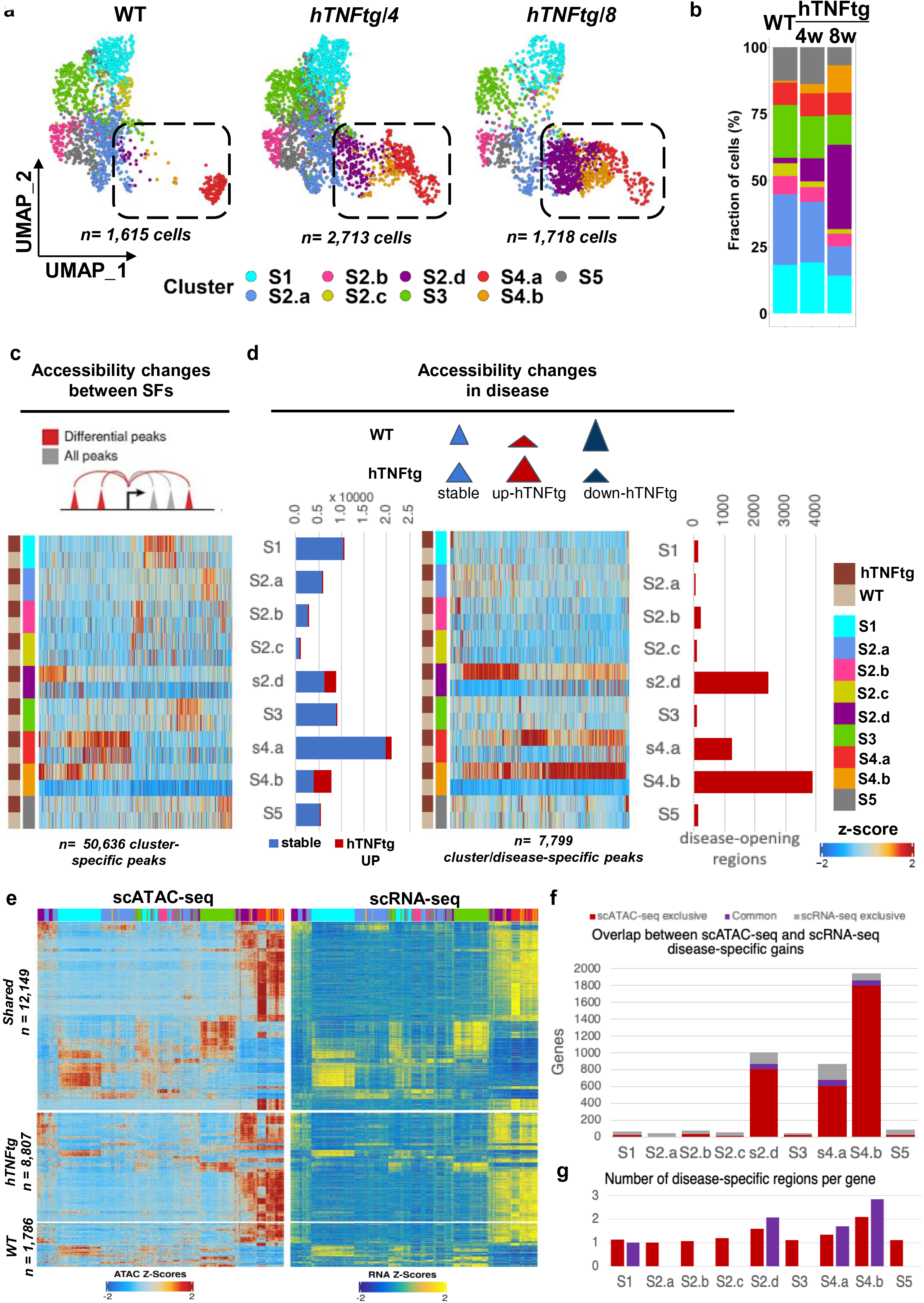
scATAC-seq recapitulates the remodeling of the synovial mesenchyme between WT healthy and hTNFtg arthritic joints. (a). UMAP representation of SFs across the three different samples (WT, hTNFtg/4 weeks and hTNFtg/ 8 weeks as indicated). Cells are colored by cluster identities and the marked area highlights the structural dynamic changes of the intermediate (I) and lining (L) subpopulations during disease progression. (b). Stacked barchart showing relative abundances (%) of clusters across samples (WT, hTNFtg/4 weeks and hTNFtg/ 8 weeks). (c). Upper panel: Schematic representation of the marker peak detection procedure. Lower panel: Heatmap showing the clustering according to z-scores of normalized accessibility for 50,636 marker peaks across SF subpopulations and disease states (WT, hTNFtg). (d). Upper panel: Schematic representation of the disease specific marker peak definition. Lower panel: left, Stacked barcharts depict the proportions of stable (shared between WT and hTNFtg) and hTNFtg-specific marker peaks; middle, Heatmap showing the clustering according to z-scores of normalized accessibility for 7,799 disease-specific marker peaks (hTNFtg-up-regulated) across SF subpopulations for WT and hTNFtg cells. Right, Barchart depicting the number of disease-specific marker peaks (disease-opening regions) per cluster. (e). Heatmap showing the z-score of normalized accessibility and integrated gene expression of 22,742 peak-to-gene links across WT and hTNFg SF subpopulations. Upper part, peak-to-gene links that are shared between disease states. Middle part, peak-to-gene links that are unique to hTNFg cells. Lower part, peak-to-gene links that are unique to WT cells. (f). Stacked barchart depicting the number of disease-specific marker genes (described in e, middle part) exclusively found in scATAC-seq data (red), shared between modalities (purple) and exclusively found in scRNA-seq data (grey). (g). Barchart depicting the number of regions per genes with gains in accessibility detected in disease for genes found only in scATAC-seq data (red) or showing combined up-regulation in scATAC-seq and scRNA-seq (purple).

To interpret the increased expression for the disease-specific genes characterized above and to determine the relevant biological effects of the global gain in RNA polymerase II activity and TFs binding sites reachability, we determined peak-to-gene linkages (Figure 3e and methods). Many gene regulatory links (genes associated with given OCRs) appeared conserved across conditions (Figure 3c) and did not display noticeable changes at the chromatin level; this finding agrees with the observation that a large majority of the OCRs remain stable (Figure 3d, left). In contrast, for the OCRs that change upon disease we reveal 1,786 and 8,807 regions/gene associations that distinguish healthy and *hTNFtg* SFs (Figure 3e). Many up-regulated genes during the disease show a parallel gain in accessibility, particularly for the intermediate cells (S2d and s4b) where up to 40% of the genes were up-regulated in *hTNFtg* SFs, according to scRNA-seq, also showed chromatin opening in at least one of the associated OCRs (figure 3f, 61 of 151 genes for cluster S4b). In fact, for the genes commonly exhibiting scRNA-seq and scATAC-seq increase, we find chromatin opening at a larger proportion of their associated regulatory regions compared to the genes that are not differentially expressed and only show chromatin opening (Figure 3g). We conclude that key pathogenesis driver genes are robustly activated when cells simultaneously open a minimal set of linked regulatory regions.

### Gene regulatory networks controlling SFs homeostatic and pathogenic functions

To determine which TF might control the cell-type or disease-specific regulatory regions and associated genes, we performed DNA motif analyses (Figure 4a, Suppl. Figure 13a). First, we highlight cluster-specific groups of TFs likely to maintain diversity in SFs function (Suppl. Figure 13) For instance, C/EBP family of TFs, involved in many processes including cell differentiation, inflammation, aging etc (discussed in^48,49^) are linked to S5 cluster while GATA family of TFs that regulate mesenchymal stem-cell differentiation transition (discussed in^50^) is linked to S2b. In contrast, S2d and S4b intermediate subpopulations are linked to Nfatc, which is known to play a central role in bone and joint remodeling during RA pathogenesis^51^ and S4a and S4b clusters are linked to a combination of TFs including the chromatin remodeling mediators Smarcc1, Bach1/2, and the pro-inflammatory effectors Junb/d, Rel and Nfkb (Figure 4a, Suppl. Figure 13a). TF binding sites (TFBS) that appear in diseased cells (within peaks found to be more accessible in *hTNFtg* SFs) revealed Rel, Nfkb, Junb/d and Runx1 TFBS (Figure 4a). We corroborated this finding by inferring the co-accessibility scores of regulatory regions modelled per-cell by employing cisTopic^52^ (Suppl. Figure 14). By analyzing the aggregated WT-*hTNFtg* space, we identified 12 topics that show distinct contribution probabilities along the SFs (Suppl. Figure 14b, c). In particular, topic 12 matches S4a subpopulation, topic 5 matches S4b subpopulation, and topic 8 matches S4b and S2d subpopulation (Suppl. Figure 14c, d) Motif analysis was applied on the regions defining these topics and confirmed that the intermediate and lining states are controlled by master regulators including Klf, Dlx, Creb3, Runx1, and Nfkb (Suppl. Figure 14e).

**Figure 4.**
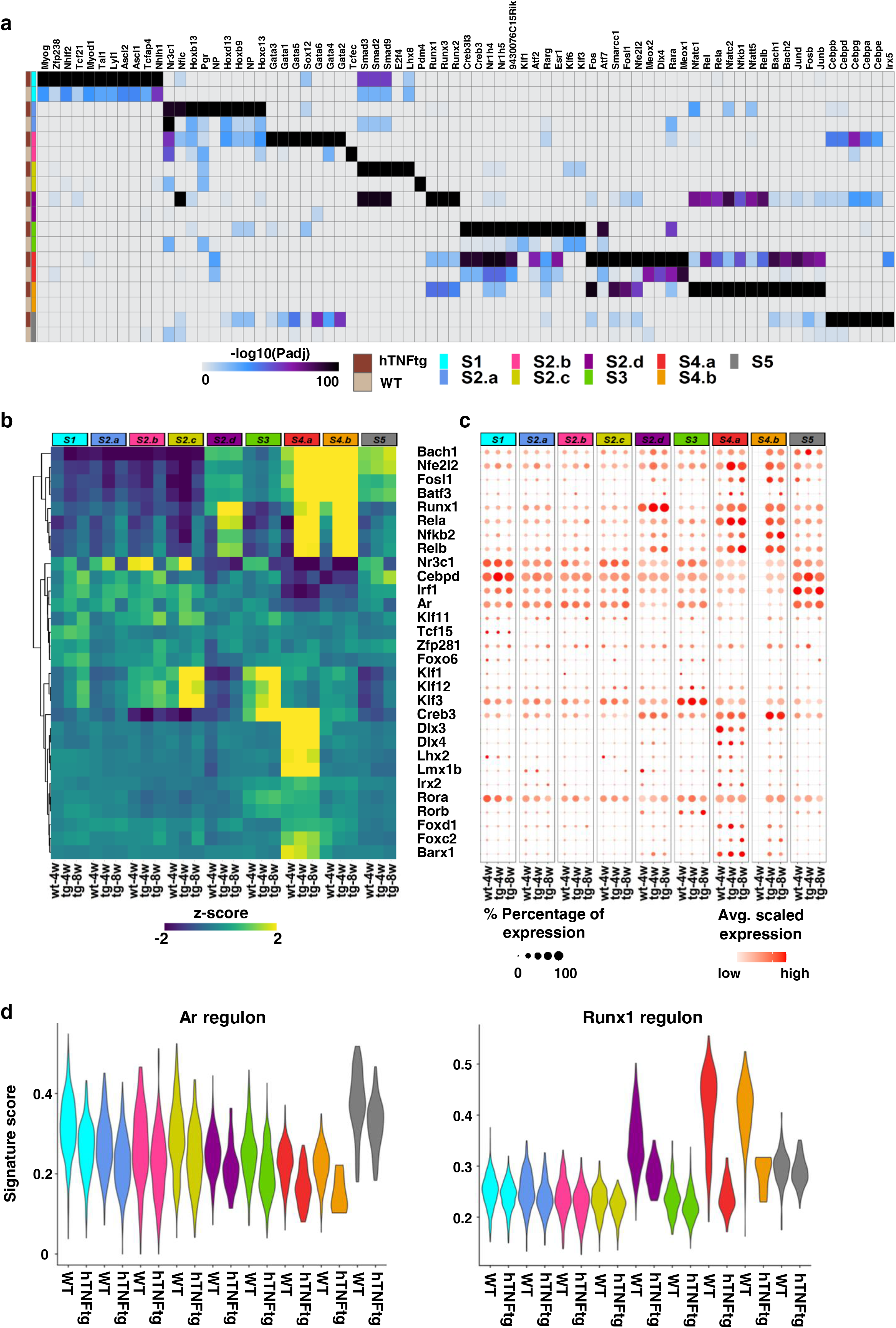
Integrative analysis of scATAC-seq and scRNA-seq infers putative arthritic regulatory programs. (a). Heatmap showing the clustering of TF motifs according to P-adjusted values (scores of motif accessibility) (see Methods) and displayed for healthy and disease state (WT, hTNFtg as indicated). Motif enrichment analysis was performed within the disease-specific marker peaks depicted in Figure 3d (right panel). Color signifies the magnitude of the enrichment (−log10 (P adjusted value), hypergeometric test). (b). Heatmap showing the average motif activity across SF subpopulations and samples (WT: wt-4w, hTNFtg/4 weeks: tg-4w and hTNFtg/ 8 weeks: tg-8w as indicated), as expressed by deviation z-scores (chromVar). Displayed regulators are all the TFs for which gene expression is positively correlated with TF motif accessibility in at least one cluster (Pearson correlation > 0.5, P-adjusted value < 0.05). (c). Expression dotplot of positive TF regulators shown in (b). The color of the dot shows the intensity of expression, and the size denotes the percentage of cells expressing the gene in each cluster and condition. (d). Violin Plots of gene signature scores across SF subpopulations and disease states (WT, hTNFtg as indicated). Gene signatures of Ar and Runx1 are composed of 23 and 183 regulated genes respectively. The target genes show significant increased expression in sublining cells (Ar) or intermediate and lining cells (Runx1) and are linked to differentially accessible peaks between disease states that are enriched with Ar or RunX1 motifs.

We resolved true “positive TF regulators” by establishing which TFs show high correlation between motif accessibility and TF mRNA expression at a single cell resolution^53^. The most deviant TFs were detected in the expanding intermediate and lining clusters (S2d, S4b, S4a) and to a lesser degree in the S5 subpopulation (Figure 4b). While we see stable high deviation scores for a subset of TF regulating the Prg4^high^ lining cluster (*Dlx, Lhx* and Lmx), we highlight notable changes in TF regulatory programs (regulons) during disease progression for the intermediate and lining cells S4a, S4b and S2d (compare healthy joint vs early and established disease states), which are operated via the TFs Nfkb, Rela, Relb, Rel and Runx1 (Figure 4b and Suppl. Figure 13b). Although, Thy1+ sublining clusters show lower deviation scores and less dynamic changes, we note that they are controlled via *Klf*, *Cebpd*, *Ar*, and Nr3c1. We validated these findings by verifying that the underlying expression scores of the TFs and of the genes they control (Gene regulatory networks-GRNs) parallels the motif deviation patterns (Figure 4c, d and Suppl. Table 5).

### A defined trajectory yields pathogenic SFs in diseased joints

We next questioned which cells give rise to the emerging S2d, S4b and S4a SF states in disease and what transcriptional transitions occur during arthritis progression. We performed cellular trajectory analysis by applying scVelo toolkit^54^ (Figure 5a and Suppl. Figure 15a) and traced the cells along an underlying Markov process to determine their respective latent time. This procedure enabled to identify plausible root cells and ending points and the most likely path bridging them (Figure 5a, Suppl. Figure 15a, b). Root properties were mainly found in the S2b state cells albeit cells in S5, S4b, S1 and S3 clusters also exhibited a root-like potential (Suppl. Figure 15b). Regardless of the origin, the cells transitioned via S2b, S2d, S4b (Figure 5a) and always ended in the area of S4a (Suppl. Figure 15b). Independent trajectory prediction methods consistently found a continuum of S2b towards S2a, S2d, S4b and S4a state and confirmed the existence of a pathogenic branch (Suppl. Figure 15c) that fits with the observed expansion of the underlying SF clusters during disease (see Figure 2b). We hypothesized that proliferative events could explain the gain in intermediate cells. Indeed, a subset of S4b cells adjacent to S4a showed activation of Cdk1 and Ccnb1 genes (Suppl. Figure 6a) and preferential expression of G2/M phase markers (Suppl. Figure 6b, c) indicating that proliferation might explain at least partially the increased abundance of the aforementioned cells in *hTNFtg* mice.

**Figure 5.**
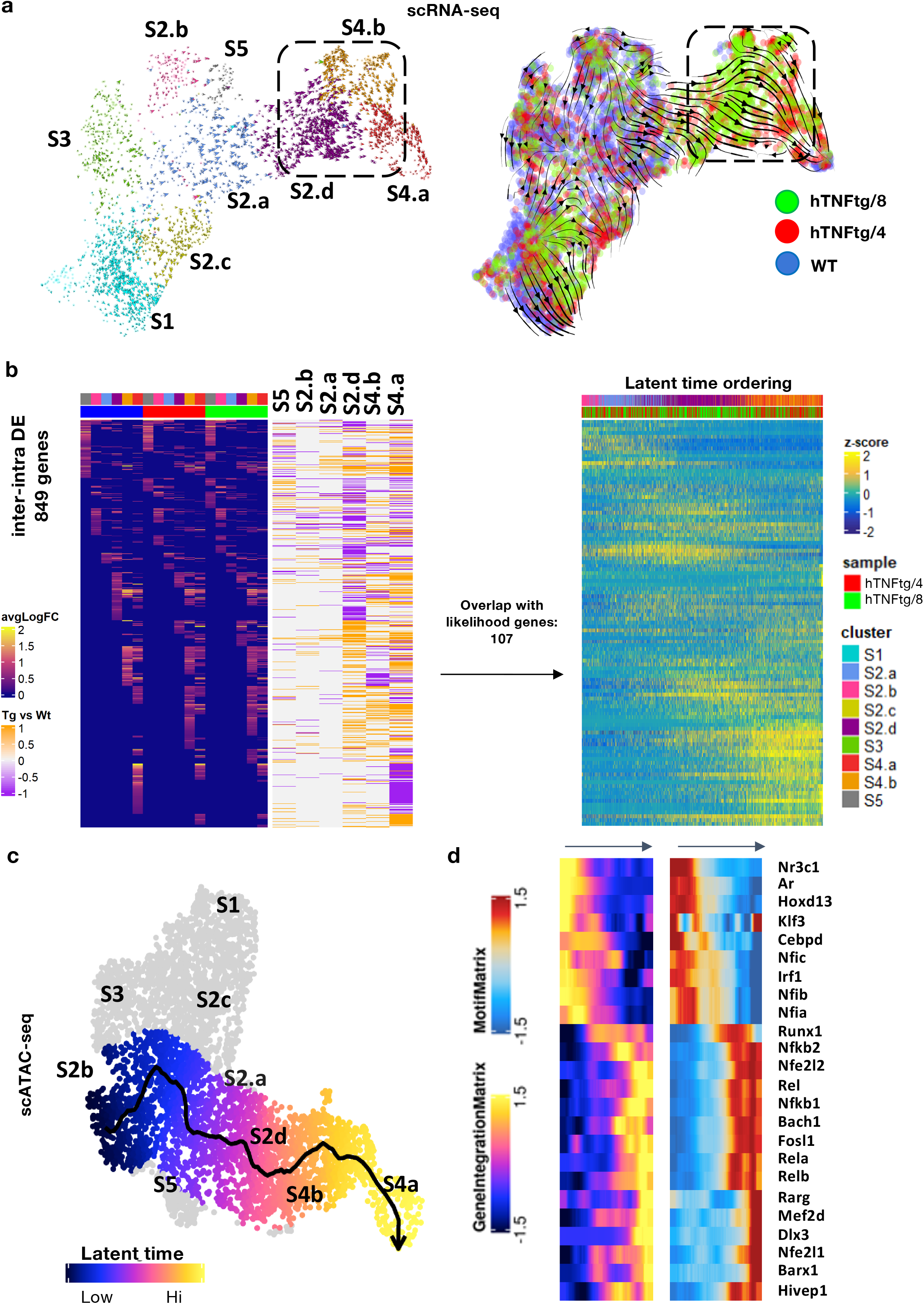
Inference of Synovial Fibroblast transcriptomic and open chromatin trajectories driving arthritis in the joints. (a). UMAP representation of RNA velocity analysis highlighting cell transitions and dynamic relations between SF clusters i. Left panel shows the large RNA velocity scores and the well-defined direction of the trajectory in the pathogenic branch (doted square) comprised of intermediate (I, S2.d and S4.b) and inflammatory lining (L, iS4.a, see Suppl. Figure 7) clusters detected in hTNFtg cells. Right panel shows the directions and amplitude of the inferred trajectory overlaid on a sample-resolved UMAP plot detailing the position of WT and hTNFtg (4, 8 weeks) samples. (b). Left: heatmap depicting the inter-cluster FC values (left) in line with the intra-cluster (Tg vs WT) FC values for the 849 genes displaying both cluster specificity (marker gene of at least one of the represented cluster) and disease-associated changes (tg vs WT) (binary values are used to denote up regulation (1, orange) or down regulation (-1, purple) of those genes in the intra-cluster Tg Vs Wt comparison, see Methods). Note the overrepresentation of genes expressed in the pathogenic branch. Right: heatmap showing the z-scores for 107 genes obtained from the inter-intra DE genes (depicted on the left) with the likelihood genes predicted by scVelo and obtained from (a). hTNFtg cells of the main trajectory (see suppl. fig 15b) were ranked according to the latent time values obtained when an S2.b cell was set as root of the trajectory. Genes were clustered according to z-score to reveal the sequence of gene expression switches observed along the process (c). scATAC-seq trajectory analysis (summarized by the black smoothed arrow) that recapitulates the existence of a pathogenic process developing through S2.d - S4.b - S4.a clusters. Color indicates the cellular fate across the inferred trajectory. (d). Heatmap showing the integrated gene expression activity (left panel) and the TF motif deviation (right panel) of the positive TF regulators described in Figure 4b along the pseudotemporal axis (left to right, black arrows) in the S2.b/S5 - S2.a - S2.d - S4.b - S4.a branch determined in (c). Note how TFs gene expression is significantly correlated with TF motif deviation across the cell trajectory.

Consistent with the TNF dependence of our murine arthritic model and the RNA velocity analysis outcome, we detected high activity scores for “response to TNF’’ for some of the suggested initial states like S2b and S5 as well as for the expanded S2d, S4a and S4b clusters (Suppl. Figure 15d, upper panel). Intriguingly, S2b exhibit transcriptional proximity to Notch3+ S5 signature (Suppl. Figure 15d, lower panel), supporting and complementing the recent evidence, which implicates Notch3 signalling in assigning the perivascular identity and priming transcriptional cues of SFs during autoimmune arthritis^20^.

To understand potential functional relationships within the inferred cellular process, we reconstructed transcriptome dynamics in light of the DE status and cells position in the proposed continuum. First, we focused on a subset of genes showing both cluster- and disease-specificity. The 849 genes isolated from 2322 inter-cluster DE genes (Suppl. Figure 11a and Suppl. Table 6) showed to be mainly affected during the transition of cells into the intermediate (S2d and S4b) and pathological-lining states (S4a) (Figure 5b). 107 genes that were also identified by scVelo as drivers of the differentiation process thanks to their high likelihood gene scoring (Suppl. Figure 11b, and Suppl. Table 6) were certified to play crucial roles in the progression of disease (GO analysis, (Suppl. Figure 11b and Suppl. Table 7)). Assignment of those genes in three main categories - early activation, intermediate activation, and late activation, based on the output of hierarchical clustering of the gene expression scores, revealed the structure of the transcriptional pattern driving cellular changes from the initial to the final state (Figure 5b) with genes such as *Runx1*, *Cd44*, *Tnfaip3* and *Tnfaip*6, *Icam1* or *Inhba* also found (Suppl. Table 6).

The cellular path was also confirmed from open chromatin data (trajectory inference from scATAC-seq^55^) and pseudo-temporal ordering of the cells recapitulated at the epigenetic level the pathogenic transitions observed with scRNA-seq (Figure 5c). We next hypothesized that the positive regulators that show increased activity along the axis S2b/S5-S4a are essential for arthritic genes regulation and we confirmed by correlation analysis between TF gene expression and the respective motif accessibility which TFs drive the differentiation during pathogenicity. Indeed, in accord with the regulon analysis above, we found that Runx1 denotes a “switch” activating the expansion and development of disease-specific S2d, S4b and S4a subpopulations and drives directly 27 of the 107 genes we defined as essential to atherogenicity (Suppl. Table 5), while TFs like Rel, Nfkb2, Dlx3, Bach1 are key effectors of this process (Figure 5d). Together these results suggest that the expansion of the S2b-S2a-S2d-S4b-S4a branch upon TNF expression commands arthritis development and influences cell fate choices via specific sets of pathogenesis induced genes.

### Common transcriptional modules control SFs in human RA and murine *hTNFtg* inflammatory arthritis

We integrated the previously generated scRNAseq data from synovial biopsies of RA patients^16,17,19^ (H), with our *hTNFtg* scRNAseq dataset (M) (see Methods for details). We found that cells of both species align particularly well in the newly defined UMAP space (Figure 6a, Suppl. Figure 16a), and unbiased graph-based clustering identified seven (H1-H7; M1-M7) sub-populations (Figure 6a, Suppl. Figure 16a). Correlation heatmap of the MVGs between human (H) and mouse (M) clusters revealed significant similarities in SFs expression programs in the two species, albeit for cluster 2 that contains mainly human SLSFs and only few mouse cells derived from the SLSFs that we described above (Figure 6b). The mouse SLSF populations S1, S2a-c, S3 and S5 located principally to clusters 3 and 4 and matched previously annotated human sublining cells expression profiles (Figure 6b, and Suppl. Figure 16-17). The cluster 1 and, to lesser extent, the cluster 7 brought together the human and murine Lining Prg4^High^ cells (Figure 6b and Suppl. Figure 17b). They also contain a previously under-appreciated proliferative mixed Lining/Sublining SF state (see below), fitting with the idea of their cellular expansion in diseased joints. Finally, Cluster 5 contains the bulk of the mouse S2d (Dkk3+) SLSFs and M5 is linked to human cells in both clusters 5 and 6, suggesting that both of these human clusters (H5 and 6) likely acquire the “intermediate” arthritis-specific profile we identified in *hTNFtg* SF states (Figure 6b, and Suppl. Figure 17b).

**Figure 6.**
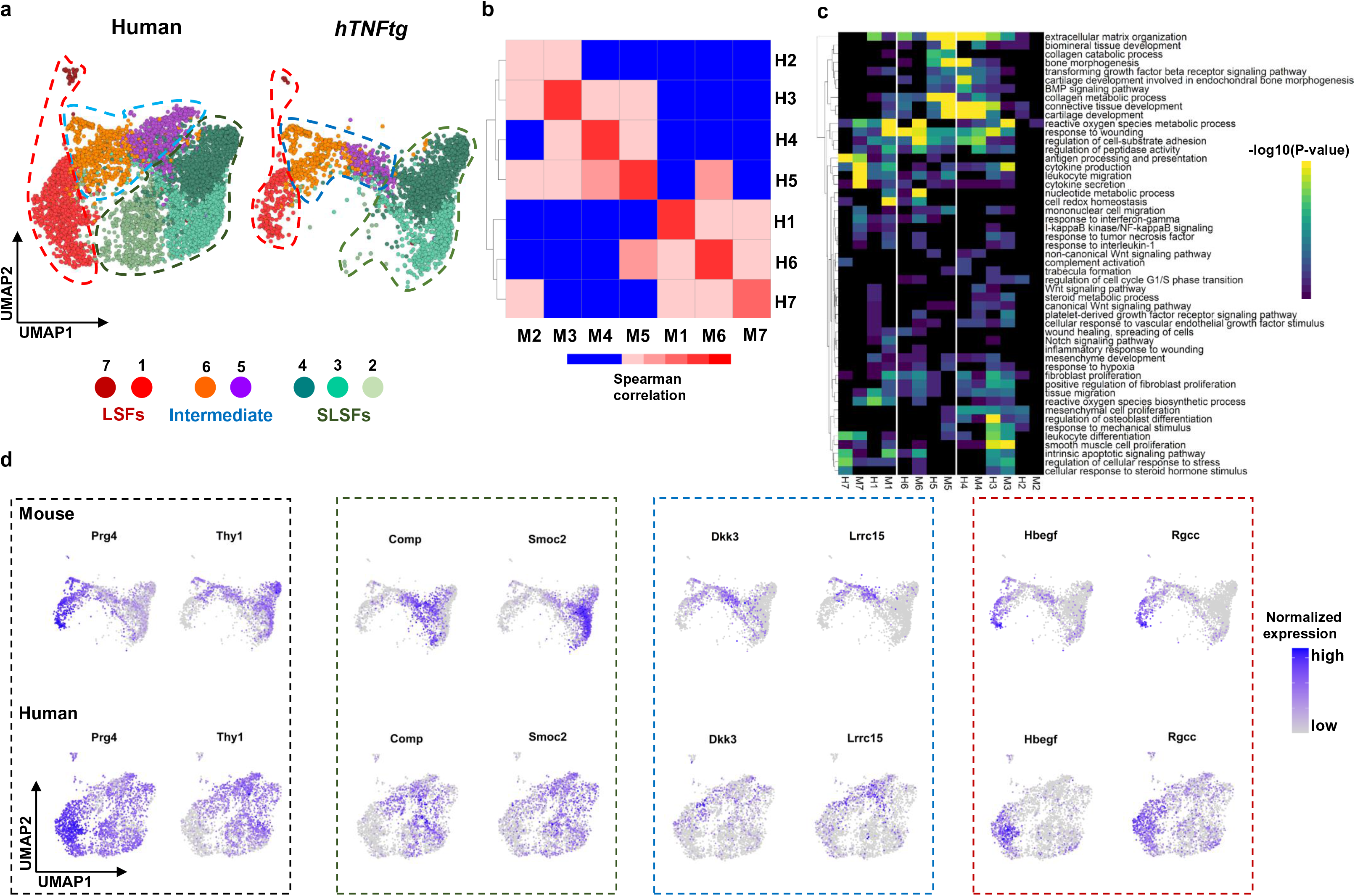
Integrative analysis of SFs from hTNFtg 197 murine model and human RA joints. (a). Integration of 24,042 RA patients’ SFs (3 different studies, Zhang et al., 2019, Wei et al., 2020 and Stephenson et al., 2018) with our 3,051 hTNFtg SFs identified 7 human(H)-mouse(M) aligned SF clusters. Separate plotting of UMAPs distribution for the pooled human (downsampled to 3051 cells) and mouse datasets cells colored by cluster identity highlights the striking inter-species overlap for homeostatic, intermediate and lining groups. (b). Correlation heatmap (average expression of MVGs) between human and mouse SF clusters. Note the highly significant similarities in H and M expression programs within a given cluster and within functional groups of cells. (c). Heatmap showing a selection of enriched functional terms and pathways in H and M clusters. Note the co-clustering of H and M in a given group of cells, and the segregation of distinct GO term enrichment for each functional group. (d). Feature plots of representative marker genes commonly expressed between homologous H and M cells for homeostatic (green box), intermediate (blue box) and lining (red box) groups, as well as for Prg4 and Thy1 genes (black box).

Functional inter-species similarities were confirmed via GO enrichment analyses of marker genes and co-clustering of (H) and (M) groups (Suppl. Tables 8 and 9). We highlight conserved functions and processes of SLSFs in regulating vasculogenesis, cell proliferation, muscle tissue development, bone and tissue renewal (clusters H3, M3, H4 and M4) (Figure 6c). We demonstrate that M5 and H5 clusters are marked by pathogenic RA features such as metalloproteinase secretion, collagen catabolic processes and bone destruction signaling pathways, further supporting the similarities with the Dkk3+ SFs in *hTNFtg* model. Clusters 1, 6 and 7, which contain SFs from the lining synovial compartment that were previously acknowledged for their destructive properties, display pro-proliferative pathways, but also appear to regulate immune-related and adhesion/migration pathways (Figure 6c). In addition, key marker genes show good levels of conservation between mouse and human data (Figure 6d).

At the regulatory level, integration of human and mouse data using SCENIC algorithm allowed the inference of common TF regulatory networks (regulons). Briefly, we first identified co-expressed genes to formulate putative regulatory links but retained only regulatory links with direct motif relationship between genes and TFs. Finally, we scored each regulon in each cell using AUC analysis (see Methods). We then preserved all the common and conserved TFs operating in datasets from the RA patients and arthritic mice. We identified the mouse regulatory modules (clusters of TFs) by applying pairwise correlation between the motif deviations of the mouse/human conserved TFs, and applied hierarchical clustering, as previously described^56^. This approach identified three main regulatory modules defining lining, intermediate and sublining states and demonstrate a substantial overlap across species (Figure 7a). Regulons are governed by Ar, Dlx3 and Runx1 TF activities (Figure 7b) and GO enrichment analysis of TF and downstream genes (Figure 7c) indicated the modules shared functionalities in both species: module one (Ar) controls multipotent functions of the main core of SLSFs; module two (Runx1) conducts functions reflecting a rather inflammatory profile consistent with the intermediate profile of our *hTNFtg* SLSFs and we find up to 25 of the 107 core mouse genes as target genes in human cells (Suppl. Table 10) highlighting the translational potential for genes like Tnfaip3 and 6, Tlr2, Lrrc15, and Bmp2. Of note, module three (Dxl3) exhibits less acknowledged functions, which should be related to the lining SF profile of human and mouse SFs (Figure 7c and Suppl. Table 10).

**Figure 7.**
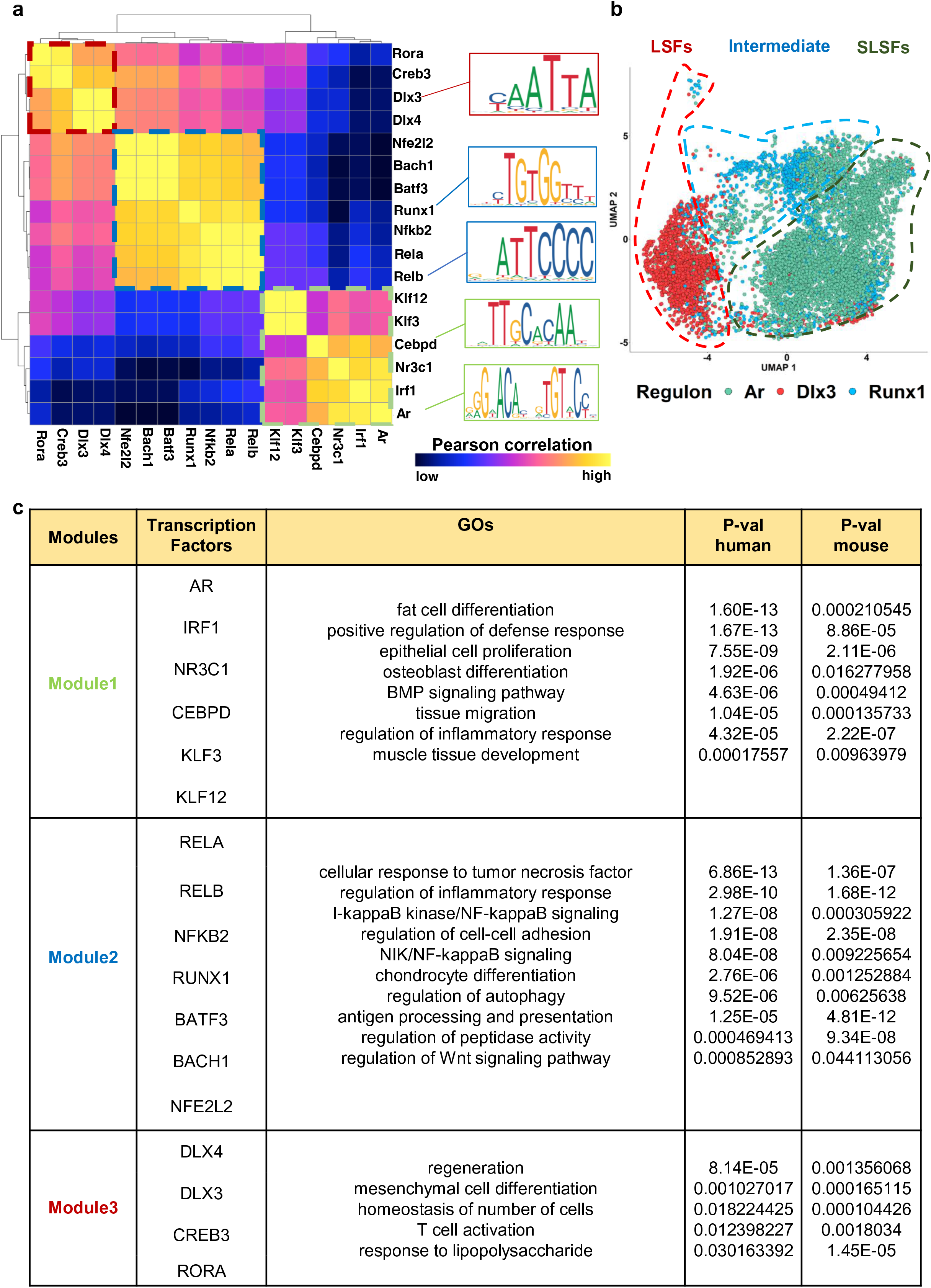
Shared Gene Regulatory Networks (GRNS) in SFs of hTNFtg mice and human RA. (a). Regulatory network analysis in mouse and human datasets reveals 17 shared regulons. Correlation and clustering analysis according to hTNFtg data reveals organization of the regulons in 3 main modules. Shown on the right are representative underlying TF motifs for each group. (b). UMAP plot depicting the cells with activity of regulons AR, RUNX1 and DLX3 for the human data. (c). Summary table of the GO enrichment analysis of the modules shown in (a). For each module we indicate the TFs (displayed in the second column) and the commonly enriched GOs and mouse and human regulons respective p-values (third to fifth column).

In conclusion, our integrative approach establishes shared mouse-human SF subsets with highly similar chromatin and transcriptional programs and functional characteristics, validating the predictive capacity of *hTNFtg* model towards deeper mechanistic understanding and effective preclinical research and development for the discovery of novel SF-targeted therapeutics.

## Discussion

In this study we postulated that defining the differences between normal and pathogenic fibroblast states in the *hTNFtg* mouse model of polyarthritis, will help us uncover and segregate the distinct fibroblast profiles and the underlying regulatory networks that specifically characterize the TNF-mediated arthritic pathology.

### The normal synovium

We report here for the first time that the normal synovium exhibits different SF states, which reflects the complexity of the SF tissue serving different functions to maintain homeostasis. Owing to their mesenchymal origin, normal SF states segregate by their responses to growth factor and differentiation signals such as WNT, BMP, TGFb. In line with the variety of elicited responses and the diversity of observed states, our GO analysis was essential to fully appreciate the related functionalities of SLSF (Thy1+) clusters regarding angiogenesis control (S3 cluster), osteogenic processes (S2 clusters), chondrogenesis and muscle development (S1 cluster). Similarly, Thy1-LSFs present a less pronounced profile, regulating lining layer size though apoptotic and migrative properties. By their specific secretome (predicted here by the transcriptomic signature), lining SFs directly respond to wounding, whereas the mechanisms to regulate mitochondrial calcium levels possibly contribute to the proper signaling alertness. Detailed mechanistic understanding of the segregation of functions in the homeostatic synovium should help to design strategies to help preserve its beneficial properties that seem perturbed during pathogenesis.

The lineage inference of SFs in homeostatic conditions indicates three major poles of initiating SFs with cells from S1 and S2b clusters keeping SLSFs character but moving mainly towards S3 and S2a/S5 states respectively and an “autonomous” S4a activity presenting a stem-cell like progenitor character for healthy lining SFs (Suppl. Figure 17). The analysis of the regulatory networks behind transcriptional SF states revealed the homeostatic TFs governing the healthy synovium, supporting the identity prediction for heathy SFs. We identified members of Dlx family (Dlx3 and 4) and Lhx (Lhx2 and 9) to be steadily expressed in healthy Prg4^high^ LSFs. Although these factors have not been yet studied in SF biology, they are known to facilitate crucial functions in the skin, in skeletal formation and development, and in tooth morphogenesis^57–59^, indicating that they may have a counterbalancing role in the regeneration of joint structures. LSFs are also characterized by the expression of *Tgif1*, which has been reported to act both as promoter or repressor of cancer, depending on the context. It is a repressor of retinoid^60^ and TGFb signaling, which also recruits histone deacetylases’ activities^61,62^, regulating chondrogenesis^63^ and inflammatory phenotype of endothelial cells^64^, suggesting a similar role in the LSF homeostatic phenotype. Along the same line of evidence, the members of GATA family regulate S2b SFs (GATA-4 and GATA-6), are well linked to mesenchymal responses towards mechanical stress in heart tissue^65^. However, their role in joint homeostasis is still poorly defined. Interestingly, GATA6 enhances the expression of smooth muscle markers such as α-SMA, SM22-α upon TGFb treatment of mesenchymal stem cells^66^, indicating that it may also affect similar responses and activation state of the GATA6-expressing SLSFs. S1 SFs are marked by Col15a1, governed by Tcf15, a transcription factor that labels the primitive identity of haemopoietic stem cell, thus supporting their role as an ancestor SF state^44,67^.

### The diseased synovium in TNF-mediated arthritis

In our comparative analyses we delineated the nature of changes in the arthritic stroma, demonstrating a relative remodeling of both sublining and lining SF molecular profiles. In light of the structural properties and homeostatic functions of the SF clusters identified in the healthy synovium, we detected a differential expansion of an inflammatory lining profile at the expense of the healthy lining SF signature (Suppl. Figure 7). Similarly, we noticed a relative reduction of the homeostatic sublining SF profiles (S1, S2a, c, S3, S5 and S4a, Figure 2) possibly reflecting aberrations in their functionality during disease, therefore strongly supporting the previously reported aberrations in reparative functions^68,69^ in RA tissue. In line with the reported expansion of the sublining SF compartment in RA^18,19^, we identified two emerging, arthritis-specific Thy1+ SF intermediate subpopulations (S2d and S4b). The S2d cells are defined by the expression of Dkk3 and Lrrc15, two markers that had been individually described and recently linked to emerged pathological states of fibroblasts in RA^19^ and other inflammatory and cancerous human conditions^44,70^ respectively. Tissue remodeling and inflammation are among the functions characterizing the expanded sublining S2d (Dkk3/Lrrc15+) SF type. The other expanded S4b SFs also express Dkk3, however, their state is mainly characterized by high proliferative and DNA imprinting capacity, indicative of the structural and epigenetic changes reported for RA^71–73^. Remarkably, the Thy1+ S4b SFs share a high Prg4 expression pattern along with other features of lining Thy1-SFs. Since these two fibroblast states are absent in healthy conditions, their expansion could be suggestive of a TNF-mediated pattern of SF differentiation in arthritic disease. This hypothesis is advocated by the lineage inference based on scVelo showing the major differentiation cue in murine diseased synovium: the sublining S2b SFs represent a root fibroblast state from which the two arthritis-specific SF subpopulations emerge. The differentiation program tends to aim towards Prg4^high^ SF state and suggesting the fate of SLSFs (Thy1+) as a continuum towards LSFs (Thy1-) in disease. This transcriptional cue is in line with the detected expansion of the inflammatory lining profile (S4a-Suppl. Figure 7) indicates the S2d and S4b states as an intermediate stage in the progressive expansion and differentiation of the arthritic SFs.

### Common regulatory networks governing murine and human diseased synovium

We evaluated the translational impact of these results by comparing our murine data with the available scRNAseq data derived from patients with RA. We identified a high degree of similarities cluster- and function-wise. Interestingly, the DKK3/LRRC15+ SFs did exhibit high expression similarities between species and in line with previous observations, they acquire an intermediate signature lying between the sublining/perivascular Notch3+ and the Prg4^high^ lining SFs in both human and murine arthritic synovium (Suppl. Figure 18 generated with data described in ref^20^). Whether this shared feature among human RA and both murine arthritic models (chronic-*hTNFtg* and acute Serum Transfer Induced Arthritis-STIA) depends on upstream, arthritogenic TNF signals directly, or indirectly through the secondary induction of Notch signaling as previously reported^74^, remains to be explored in vivo. Importantly, the need for understanding if and how this intermediate SF state drives the pathogenicity and leads the destructive nature of arthritis through activation and expansion of LSFs, signifies the necessity for future targeted functional and cell-fate mapping studies.

GRNs revealed in both mouse and human datasets and the identification of the open chromatin regions (scATAC) around target genes in arthritis, enable us to uncouple the modulators of the pathogenic expansion of SLSFs to LSFs and identify unique patterns of activity. Among these highly significant factors, the NF-κB pathway components NF-κB1/2, RelA and RelB are well known to heavily regulate inflammatory processes including the inflammatory arthritic diseases^75,76^. Interestingly, we have previously addressed the SF-specific NF-κB mediated responses in the development of arthritis of the *hTNFtg* mice and we mechanistically showed that a major NF-κB activator, the IKK2 kinase, acts as dual modulator of arthritis though both the inflammatory and the death responses of SFs^6^. Oxidative stress regulator Nfe2l2/NRF2 is highly expressed during disease, along with factors such as Bach1, Fosl1 and Mef2d, which have been shown to control NRF2 activity^77–79^, suggesting a molecular network related to the extensive oxidative stress heavily reported for RA^80^, and highlighting specific components of a pathway with therapeutic potential in TNF-mediated arthritis. Interestingly, Nrf2/Keap1 pathway can inactivate NF-κB activity through ubiquitin-mediated degradation of IKK2^81^, suggesting links between the deregulated NF-κB and NRF2 pathways in arthritis. Additionally to hematopoiesis, Runx1 has been associated with the osteochondral differentiation (along with Runx2 and 3), the myofibroblastic activation^47^ and it has been suggested as a dual inflammatory modulator and even an epigenetic modifier, depending on the context^82–88^. Owing to its robust upregulated expression, Runx1 emerged as an essential master regulator of DKK3/LRRC15+ SFs in both species, indicating a promising disease-paramount pathway that requires further studies. Conclusively, the integrative analysis showed that the *hTNFtg* SFs shares many commonalities with human RASFs both at transcriptional and epigenomic levels, suggesting an added predictive value of the *hTNFtg* model for integrative translational studies of SF functions.

## Conclusion

To date, this is the first study exhibiting and comparing the homeostatic heterogeneity of SFs with SFs in a diseased environment. We provide a map of transcriptomic and epigenomic profiles that allowed us to derive general principles by which sets of specific TFs and GRNs cooperate to rewire SFs identity and functions during onset and progression of arthritis. Finally, we aligned our findings with the human context to highlight the advantages of *hTNFtg* mouse model to study potential novel fibroblast-targeted diagnostic and therapeutic opportunities for RA.

## Materials and Methods

### Mice/ethical compliance

All mice were bred and maintained on C57BL/6J genetic background in the animal facilities of the BSRC Alexander Fleming under specific pathogen-free conditions in accordance with the guidance of the IACUC of BSRC “Alexander Fleming’’ and in conjunction with the Veterinary Service Management of the Hellenic Republic Prefecture of Attika/Greece. Experiments were monitored and reviewed throughout its duration by the respective Animal Welfare Body for compliance with the permission granted.

### Flow cytometry and fluorescence-activated cell sorting

Isolation of SFs was performed from both hind paws. Ankle joints were dissected from 3 WT mice at the age of 4 weeks and 6 *hTNFtg*, 3 at the age of 4 weeks and 3 at the age of 8 weeks. The tissues were disaggregated by incubation for 30 min at 37°C in an enzymatic digestion medium consisting of DMEM, 10%heat-inactivated FBS, collagenase (0.5mg ml^-1^) from *Clostridium histolyticum* (Sigma, C5138) and 0.03 mg ml^-1^ DNase (Sigma, 9003-98-9). Upon washing the cells with PBS containing DNase, they were blocked in 1% BSA in PBS and Fc blocker (unlabeled anti-CD16/32, Biolegend 101302) for 10 min at 4°C and stained with fluorophore conjugated antibodies for 20 min at 4°C (anti-Pdpn PE-Cy7, Biolegend 127411; anti-Thy1 A647, Biolegend 105318; anti-CD31 APC/Fire 750, Biolegend 102433; anti-CD45 APC-Cy7, Biolegend 103116; anti-Ter119 APC-A780, eBioscience 47-5921-80). After washing with PBS, cells were resuspended in FACS buffer (PBS, 1%BSA). Sorting of cells was performed with BD FACSAria III and the BD FACSDiva software and dead cells were excluded by Dapi staining. Sorting gaiting for Single-cell RNAseq and bulk RNAseq were different (Sup. Figure1B). For sorted populations, purity and viability, was determined by reanalysis for the target population based on cell surface markers immediately post sorting. Purity was >99% for each target population.

### Histopathology and immunofluorescence

Histological H&E staining was performed on paraffin ankle joint sections as previously described^7^. For Immunofluorescence, cryosections were probed with antibodies against Thy1 (Alexa Fluor 488 anti-mouse CD90.2 Antibody, Biolegend 105315, or Alexa Fluor 647 anti-mouse CD90.2 Antibody, Biolegend 105318, both Clone, 30-H12), Clu (Polyclonal Rabbit antiHuman CLU / Clusterin, LS-C331486, LSBio), Gdf10 (GDF10 Polyclonal Antibody, BS-5720R, Bioss Antibodies), CD31 (APC Rat Anti-Mouse CD31, 551262, BD Biosciences, clone MEC 13.3), Notch3 (anti-Notch3 antibody, ab23426, abcam), Comp (Anti-COMP/Cartilage oligomeric matrix protein antibody, ab231977, abcam), CD44 (FITC Rat Anti-mouse CD44, 553133, BD Biosciences, clone IM7), and Prg4/Lubricin (Anti-Lubricin/MSF antibody, ab28484, abcam). Alexa-Fluor 647–conjugated secondary antibodies (anti-rabbit, A21244, 1834794; anti-rat, A21247, 1719171; anti-mouse, A21235, 1868116; and anti-hamster, A21451, 1572558, Invitrogen), biotinylated secondary antibodies (anti-rat, BA-9400, and anti-rabbit, BA-1000). Images were acquired with a TCS SP8X White Light Laser confocal microscope (Leica) and with an Eclipse E800 (Nikon) microscope equipped with a Dxm1200F camera (Nikon). Imaging analysis and quantifications were performed with ImageJ/Fiji software (NIH).

### Generation of droplet-based single cell RNA sequencing data

Sorted live Pdpn^+^ CD45^-^ CD31^-^ Ter119^-^ synovial cells of hind paws of WT mice at the age of 4 weeks and *hTNFtg* mice at 2 different stages of the disease, early at 4 weeks and established at 8 weeks old mice were subjected to 10X Chromium Single Cell 3’ Solution v3. The platform was used to generate targeted 3,000 single-cell gel bead-emulsion per sample, loaded with an initial cell viability of 80%. The scRNA-seq libraries were prepared following the 10X Genomics user guide (Single Cell 3’ v3 reagent kits). After encapsulation, emulsions were transferred to a thermal cycler for RT. cDNA was purified and amplified with primers provided in the Single Cell 3’ reagents (10X Genomics). After purification with 0.6X SPRIselect beads (Beckman Coulter) cDNA quality and yield was evaluated using an Agilent Bionalyzer 2100. Using the provided enzyme fragmentation mix the libraries were fragmented, end-repaired and A-tailed. Products were cleaned using SPRIselect beads and adaptors provided in the kit were ligated. After cleaning ligation products, libraries were amplified and indexed with unique sample index i7 trough PCR amplification. Final libraries were double sided cleaned and their quality and size was evaluated using an Agilent Bioanalyzer 2100. Libraries were sequenced by pooling them in 1 lane on Illumina NextSeq 500 sequencer to a depth of 100 million reads each (one lane 75PE). The forward read included 28 bp for the 10X Barcode-UMI, followed by 8 bp i7 index (sample index), and 10 bp on the reverse read. Reads were converted to .fastq format using mKfast from cellranger v3 (10X genomics). Reads were then aligned to the mouse reference genome (mm10, Ensembl annotation release 91). The steps of read alignment, UMI counting and aggregation of individual sample count matrices into a pooled single matrix was performed using the 10x Genomics Cell Ranger pipeline(v3).

### Computational analysis of single-cell RNA sequencing data

DoubletFinder^89^ and Seurat R packages^90,91^ were used for doublet detection and quality control of the cells. Cells containing less than 500 genes or more than 10% of reads mapped to mitochondrial genome were excluded from further analysis. Downstream analysis of the data was performed using the functions of the Seurat package as described below. Normalization of the data was performed with the function NormalizeData using “LogNormalise” as the normalization method and 10000 as the scaling factor. To identify the most variable genes the function FindVariableFeatures was applied with mean.var.plot (mvp) as a selection method and the rest of the parameters set to default. Scaling of the gene expression values was achieved by the scaleData function. Principal Component Analysis on scaled values of most highly variable genes, as identified in previous steps, was performed by the function runPCA. To find the optimal number of principal components to be used during the step of clustering and non-linear dimensionality reduction, SVD k-fold cross-validation was performed with dismo R library (https://cran.r-project.org/web/packages/dismo/index.html). For the clustering of the cells a graph-based clustering approach was followed encompassing the construction of a k-nearest neighbor graph of the cells and the utilization of Louvain community detection algorithm. The functions FindNeighbors and FindClusters were used to achieve that, the first with the parameter dims set to the range 1:25 and the second with the parameter resolution set to 0.6. tSNE and UMAP non-linear dimensionality reduction methods were used for cell visualization in 2D through the functions runTSNE and runUmap using the optimal number of PCs=25. For the identification of cluster marker genes, differential expression analysis (Wilcoxon rank sum test, adjusted p-value based on bonferroni correction using all features in the dataset, group1=cells belonging to the tested cluster, group2=rest of the cells) was performed with the function FindAllMarkers excluding genes that exhibited less than 25% of expression in both cell groups or an absolute value of average log fold change less than 0.25. The same approach was followed in both pooled and individual sample analysis (in this analysis only cells belonging to the analyzed sample were used). Gene set overrepresentation analysis was conducted using the R package clusterProfiler^92^. The lists of up-regulated genes from each cluster (pval < 0.01 and avgLFC >= 0.25), as identified in the previous step, were used as an input gene list. All the active genes of the dataset were considered as the background set of genes. ‘Biological processes’ gene sets were used and obtained from the GO database. Enriched GO terms were considered those that showed an adjusted p-value < 0.05 and a gene count >= 3.

### Trajectory analysis

Velocity analysis was conducted by using velocyto v.0.17^93^ and scVelo v.0.2.3^54^. In particular, to count spliced and unspliced reads for each sample, the 10x velocyto pipeline was run in the filtered cellranger-generated BAM files, while for single-cell RNA velocity inference, the dynamical model of scVelo was applied. To predict the root and terminal states of the underlying Markov process, the respective scVelo functions were applied. The resulting root cells were used to infer the latent time ordering of the *hTNFtg* cells.

Following the results of RNA velocity analysis, the R package Slingshot^94^ and python package PAGA^95^ were utilized. During the run of Slingshot UMAP coordinates were used for the cells, while clusters S2b and S5 were set as possible starting points. The produced minimum spanning tree supported the existence of a pathogenic branch comprised of S2a, S2d, S4b and S4a.

### Human scRNA-seq Gene Regulatory Network (GRN) inference

To infer GRNs from the human integrated scRNA-seq data, the SCENIC^96^ workflow was applied in the normalized expression matrix. Briefly, initially co-expressed genes were grouped using arboreto python tool^97,98^. Next, using CisTarget^99^ all the inferred groups that included a Transcription Factor (TF) were considered as GRNs, while all genes with motif evidence of the respective TF in their regulatory space (hg38 refseq-r80 500bp_up_and_100bp_down_tss.mc9nr, hg38 refseq r80 10kb_up_and_down_tss.mc9nr.feather) were considered as valid TF targets. Finally, each formed regulon was scored in each cell, using AUCell^96^.

### Integration of human data

For the integration of human data 3 different datasets were used^16,17,19^. During the first step of the analysis, human genes were converted into mouse homologs using the Ensembl Biomart and MGI database, leading to the final set of 17,594 homologous pairs. Regarding the cells that were used, from the mouse dataset only the cells originating from pooled *hTNFtg* samples (3,051 cells) were processed and from the three human datasets, only the cells originating from RA patients (24,042 cells). After that the integration strategy described in ^91^ was followed through the Seurat package. More specifically all four datasets were processed for normalization and detection of most variable genes using the function normalizeData with default settings and FindVariableFeatures (method set to vst and number of variable features to 2,000). Anchors between all datasets were identified using the function FindIntegrationAnchors with dimensions parameter set to 30 and then these anchors were utilized to integrate the four datasets together using the function IntegrateData. The final object containing all the cells was processed in a standard way, performing the steps of dimensionality reduction, clustering and differential expression analysis. The integrated clusters were defined after using the FindClusters function with a 0.3 resolution. Finally, differential expression analysis was performed using findAllMarkers function with the following thresholds: pval < 0.01 & avgLFC >= 0.25. As regards the functional enrichment analysis, the up-regulated genes of human and mouse datasets were used as an input for Metascape^100^, significant terms and pathways (pval < 0.05) were used to assess similarities and differences across the datasets. (For all the comparisons between human and mouse described above, the final integrated object was split into two, one containing all human cells from the three different datasets and another containing all mouse cells from pooled *hTNFtg* samples.)

### Isolation of RNA and bulk 3’ RNA sequencing

RNA was isolated from sorted ankle joints synovial fibroblasts (sublining and lining) of healthy (4 weeks of age) and *hTNFtg* mice (4 & 8 weeks of age) using the RNeasy mini or micro kit (QIAGEN), according to the manufacturer’s instructions. The quantity and quality of RNA samples were analyzed using Agilent RNA 6000 Nano kit with the bioanalyzer from Agilent. RNA samples with RNA Integrity Number (RIN) > 7 were used for library construction using the 3′ mRNA-Seq Library Prep Kit Protocol for Ion Torrent (QuantSeq-LEXOGEN™) according to manufacturer’s instructions. DNA High Sensitivity Kit in the bioanalyzer was used to assess the quantity and quality of libraries, according to the manufacturer’s instructions (Agilent). Libraries were then pooled and templated using the Ion PI™ IC 200 Kit (ThermoFisher Scientific) on an Ion Proton Chef Instrument or Ion One Touch System. Sequencing was performed using the Ion PI™ Sequencing 200 V3 Kit and Ion Proton PI™ V2 chips (ThermoFisher Scientific) on an Ion ProtonTM System, according to the manufacturer’s instructions.

### Computational analysis of bulk RNA sequencing data

Quality of the FASTQ files was assessed with fastqc software (Andrews, S. (2010). FastQC: A Quality Control Tool for High Throughput Sequence Data [Online]. Available online at: http://www.bioinformatics.babraham.ac.uk/projects/fastqc/). Alignment of the reads to the mm10 genome was performed with Hisat2 aligner. FeatureCounts^101^ was utilized for the step of read summarization at the gene level. Differential expression analysis was conducted by DESeq2^102^. For each contrast differentially expressed genes definition was based on the following thresholds |Log2FC| > 0.58 & Pvalue < 0.05.

### Single cell ATAC-seq (10x Genomics)

Single-cell assay for transposase-accessible chromatin using sequencing (scATAC-seq) protocol was performed according to 10x Genomics instructions. Briefly, after sorting of synovial fibroblasts (see scRNA-seq protocol for details) and nuclei isolation, nuclei were resuspended in 1x Diluted Nuclei Buffer (10x Genomics). About 4600 nuclei were added in each transposition reaction, aiming a targeted nuclei recovery of 3000 nuclei. Transposition was performed at 37℃ for 60 mins. Generation of Gel beads in EMulsions (GEMs) using Chromium Controller (10x Genomics), was followed by GEMs incubation and cleanup, based on 10x Genomics recommendations. Amplification of libraries was performed in a Veriti Thermal Cycler (Thermo Fisher) programmed at 98 ℃ for 45 s followed by 12 cycles of (98 ℃ for 15 s, 67 ℃ for 30 s, 72 ℃ for 20 s), 72 ℃ for 1 min and hold at 4 ℃. In turn, libraries were double sided size selected using SPRI select reagent (Beckman Coulter) according to 10x Genomics recommendations. Before multiplexing, libraries were assayed on Bioanalyzer High Sensitivity DNA ChIP (Agilent), for quality check and determination of fragment size. Quantification of libraries was performed using Qubit dsDNA HS Assay Kit (Thermo Fisher, Cat. No Q32851). Next generation sequencing was performed at EMBL-Genecore (Heidelberg), using the NextSeq 500 platform for paired-end 75 bp reads.

### Computational analysis of scATAC-seq

The analysis of scATAC-seq datasets was conducted by using the ArchR suite^55^. Consequently, epigenetic maps of sorted SFs nuclei were obtained for 6,679 single nuclei. Latent Semantic Indexing (LSI)^53,103^, Louvain clustering, and UMAP dimensionality reduction was applied as described above (see “Computational analysis of single-cell RNA sequencing data”). To assign scATAC-seq cluster identity, gene-activity scores and scRNA-seq gene expression were directly aligned between the two modalities^91^, by first applying an unconstrained integration to gain prior cluster identity knowledge, that was in turn used as a guide for a more refined constrained integration^55^. This procedure grouped cells to 5 major clusters, corresponding to the previously annotated cell types described above (synovial fibroblasts, osteoblasts, chondrocytes, myoblasts/myocytes and vascular cells, Suppl. figure 3). All non-fibroblast cells were excluded from the rest of the analysis, resulting in a total of 6,046 SF cells that were re-analyzed in the same fashion. The integration process between scATAC-seq and scRNA-seq SFs labeled the scATAC-seq cells according to 9 SF subpopulations (see above), that were visualized in UMAP space (Figure 1; Suppl. Figure 3). To identify a robust merged peak set along the SF subpopulations, MACS2^104^ was applied at two separate pseudo-bulk replicates^55^. Next, iterative overlap peak-merging^105^ was applied at the level of the pseudo-bulk replicates (per subpopulation), and subsequently at the level of SF subpopulations across the whole dataset, to form a single merged peak set of 158,713 regions with a fixed length of 500 bp. In turn, peaks were annotated according to their respective genomic position (promoter, intronic, exonic, distal). Using the unified peak set, differential accessibility analysis between cells was performed to identify cluster-specific and condition-specific marker peaks (|Log2FC| > 0.58 & Pvalue < 0.01). Marker peaks were further analyzed using motif enrichment analysis ( CIS-BP database), to gain cluster-specific and sample-specific marker motifs ( |Log2FC| > 0.58 & Pvalue < 0.05). To further gain enriched motifs in single-cell resolution, chromVar analysis was conducted^106^. Consequently, to identify “positive TF regulators” in SF subpopulations, TF motif accessibility was correlated with integrated TF gene expression across cells, keeping all TFs with Pearson r^2^ > 0.5 and P Adjusted value< 0.05, resulting in 30 positive regulators. Finally, to identify the underlying GRNs, peak to gene linkages were called using correlation analysis between enhancer peak accessibility and integrated gene expression (see addPeak2GeneLinks() function of ArchR R package^55^. All links between genes and accessible regions with an annotated TF motif were marked as putative regulatory links between the respective TF and gene. Subsequently, all putative regulatory links were filtered to only keep genes that are upregulated in *hTNFtg* samples, as also peaks with increased accessibility in the disease samples.

## Acknowledgments

We thank Vaggelis Harokopos from the Genomics Facility at BSRC Fleming for performing RNA-seq library preparation and sequencing, Sofia Grammenoudi from the Flow Cytometry Facility at BSRC Fleming for sorting experiments. We also thank Konstantinos Lilakos (AntiSel, Greece) and Dimitris Karamitros for providing critical reagents and advice. The authors also wish to thank the InfrafrontierGR infrastructure (co-financed by the ERDF and NSRF 2007-2013) for excellent mouse hosting and flow cytometry facilities. Graphics was generated using Biorender.com.

---

**Supplementary Figure 1.**
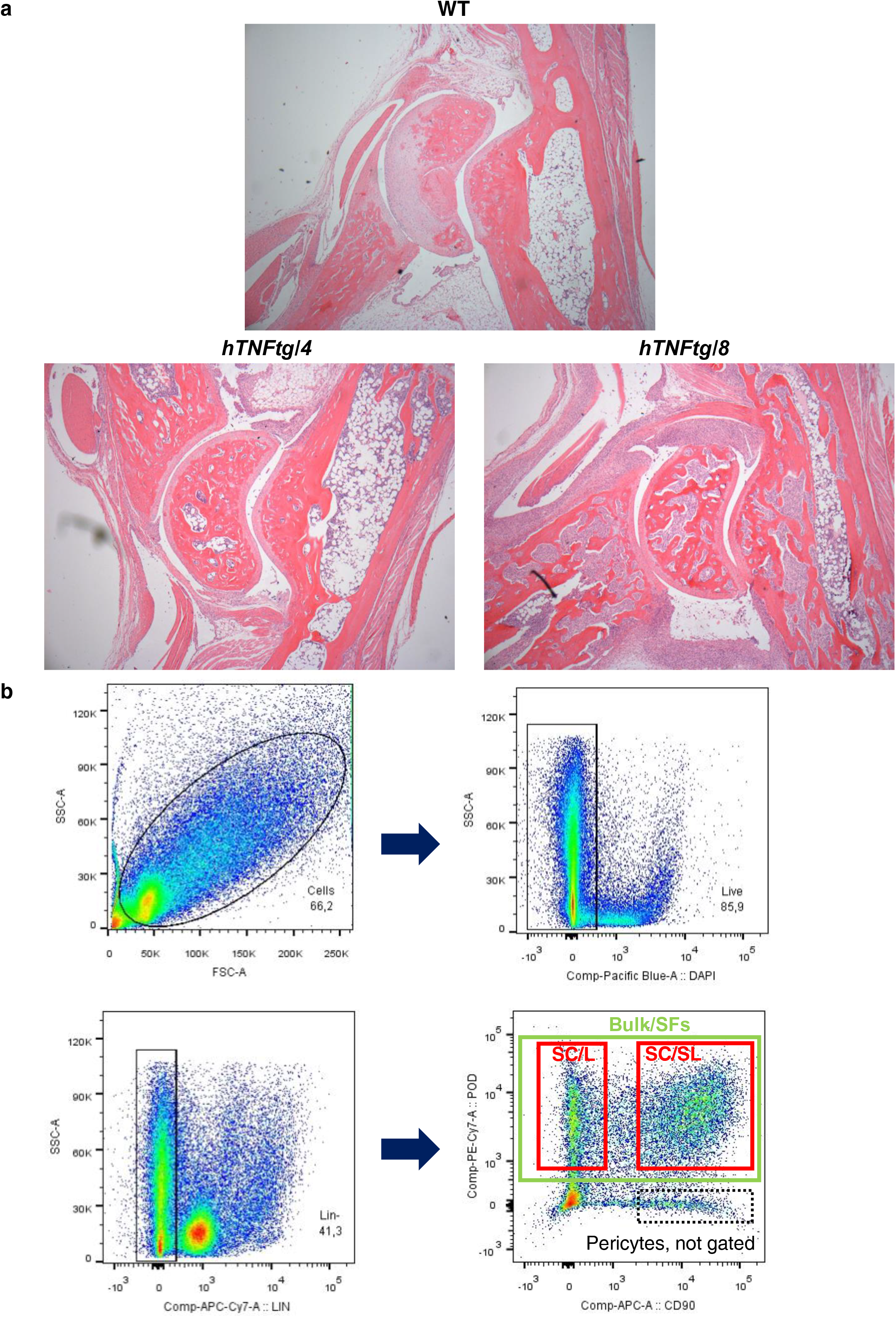
Arthritic manifestations in the ankle joint samples and sorting strategy of the digested joints employed for the scRNA-seq analysis of non-hematopoietic stromal cells of the hTNFtg and WT mice. (a). Representative haematoxylin and eosin staining of the ankle joint at the talus level from 4-week-old WT, 4- and 8-week-old hTNFtg mice (b). Gating strategy for flow-cytometric analysis of synovial fibroblasts. Intact cells are identified based on forward scatter (FSC-A) and side scatter (SSC-A) characteristics. Dead cells (Dapi), leukocytes (Cd45+), endothelial (Cd31+) and pro-erythroblasts (Ter119+) were excluded from analysis. Within the Lin- (CD45-CD31-Ter119-) gate, synovial lining (PDPN+, Thy1-) fibroblasts could be distinguished from sublining fibroblasts (PDPN+, Thy1+). The cell sorting strategy for sc analyses is highlighted with the red gating, whereas the cell sorting strategy for Bulk RNA-seq is highlighted with the green gating. Pericytes that haven’t been selected in any of the approaches are highlighted in the black doted (Pdpn-, Thy1+).

**Supplementary Figure 2.**
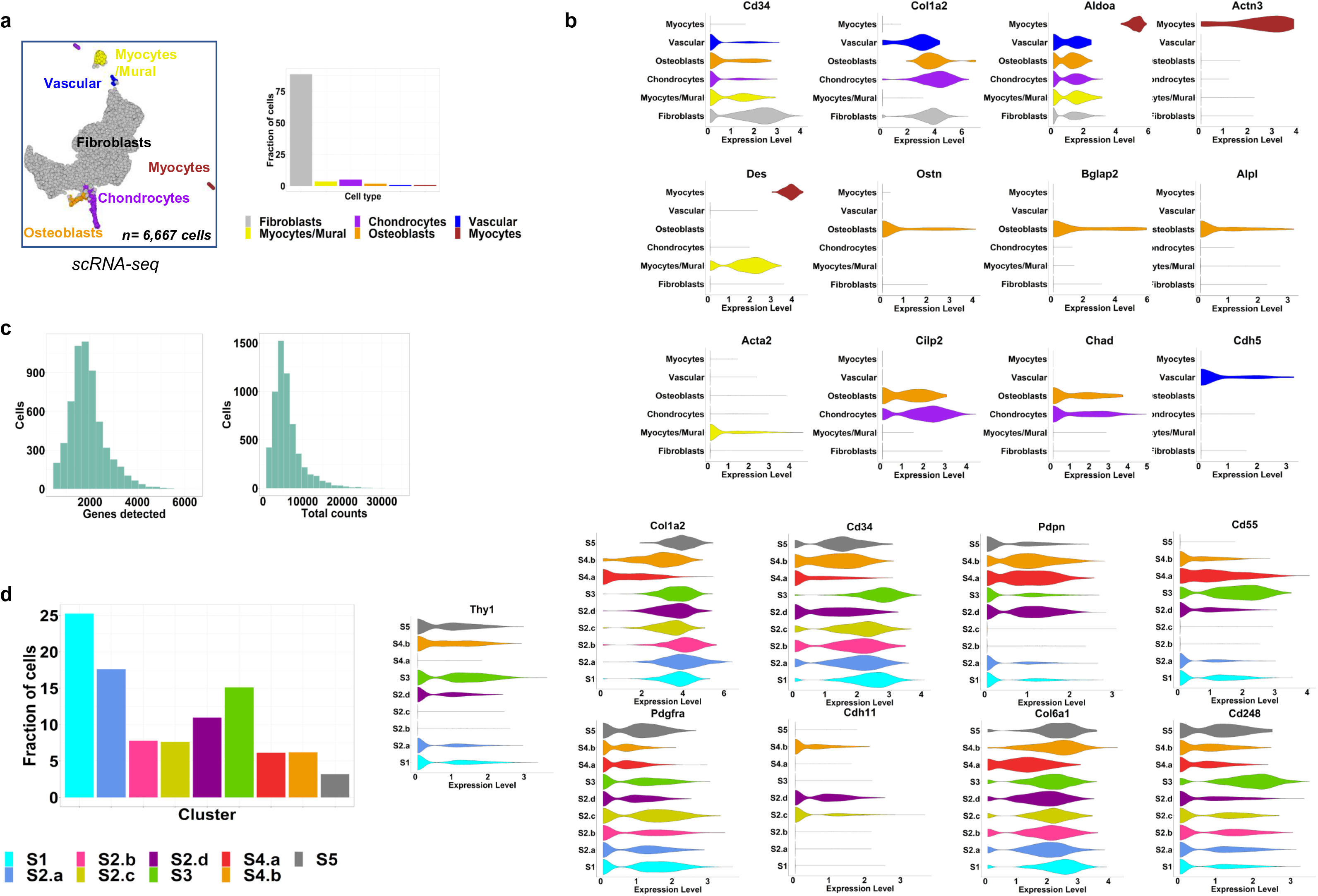
Complementary scRNA-seq analyses of non-hematopoietic stromal cells derived from hTNFtg and WT ankle synovia. (a). UMAP projection showing the different cell types in the pooled RNA-seq dataset (left) and barplot showing their relative abundance (right). (b). Violin plots showing normalized expression values of genes across the different cell types. (pooled dataset - all cells included) (c). Barplots depicting the number of genes detected per cell (left) and the number of total reads per cell (right) (pooled dataset – only fibroblast cells included) (d). Barplots depicting the relative abundances of clusters in the pooled RNA-seq data containing only fibroblasts (left panel) and violin plots showing normalized expression values for fibroblast markers. (pooled dataset – only fibroblast cells included)

**Supplementary Figure 3.**
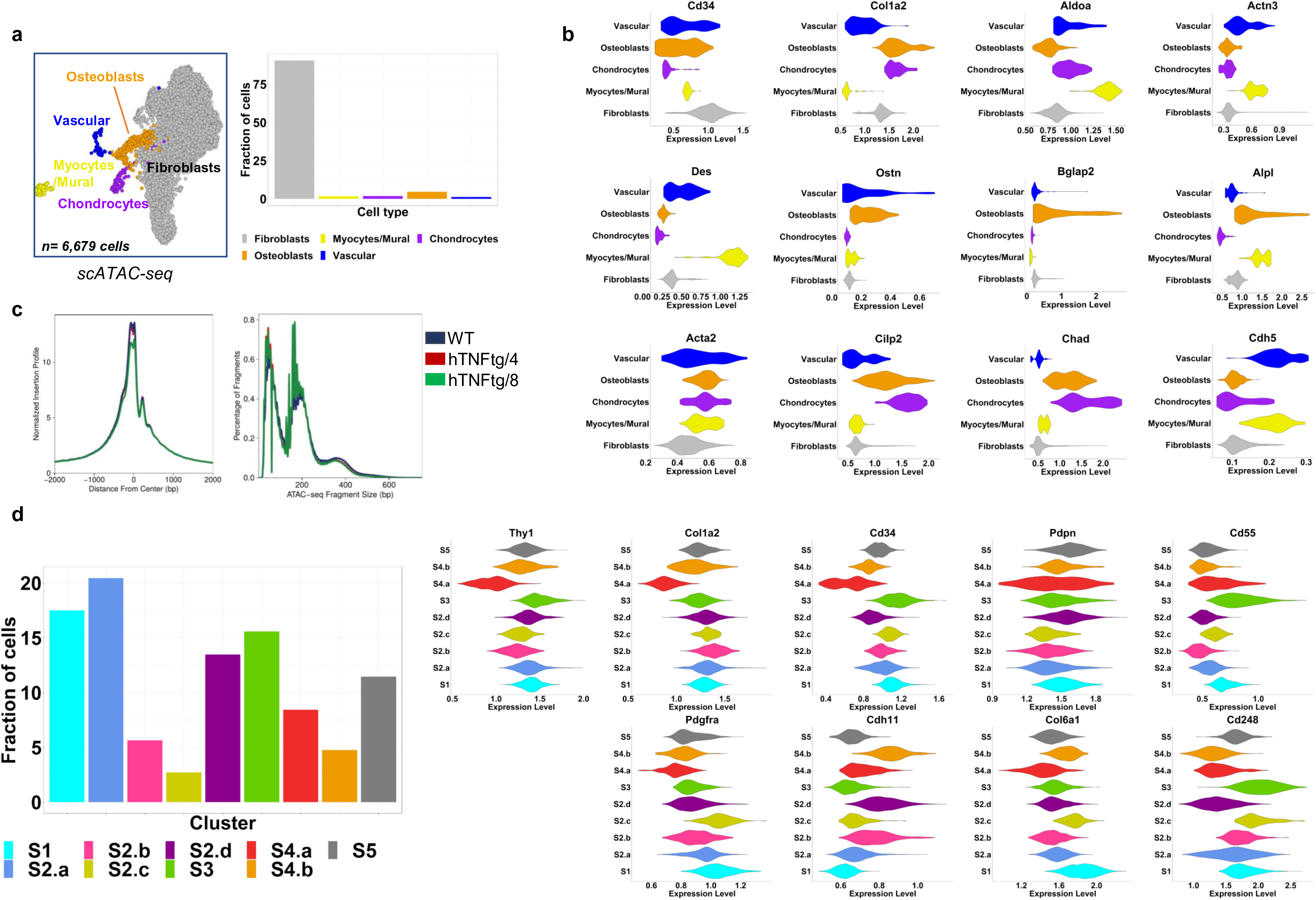
Complementary scATAC-seq analyses of non-hematopoietic stromal cells derived from hTNFtg and WT ankle synovia. (a). UMAP projection showing the different cell types in the pooled ATAC dataset (left) and barplot showing the percentage of the different cell types (right). (b). Violin plots showing gene activity scores for selected genes across the different cell types. (pooled dataset - all cells included) (c). Left panel: Average plots of reads distribution around TSSs for WT, hTNFtg/4 weeks and hTNFtg/ 8 weeks samples showing a sharp peak at TSS and a smaller peak (right of TSS) due to the stably positioned +1 nucleosome. Right panel: Plot showing fragment size distributions with negligible variability across samples (WT, hTNFtg/4 weeks and hTNFtg/ 8 weeks as indicated). (d). Barplots depicting the percentage of clusters in the pooled ATAC-seq data containing only fibroblasts (left panel) and violin plots showing gene activity scores for fibroblast markers. (pooled dataset – only fibroblast cells included)

**Supplementary Figure 4.**
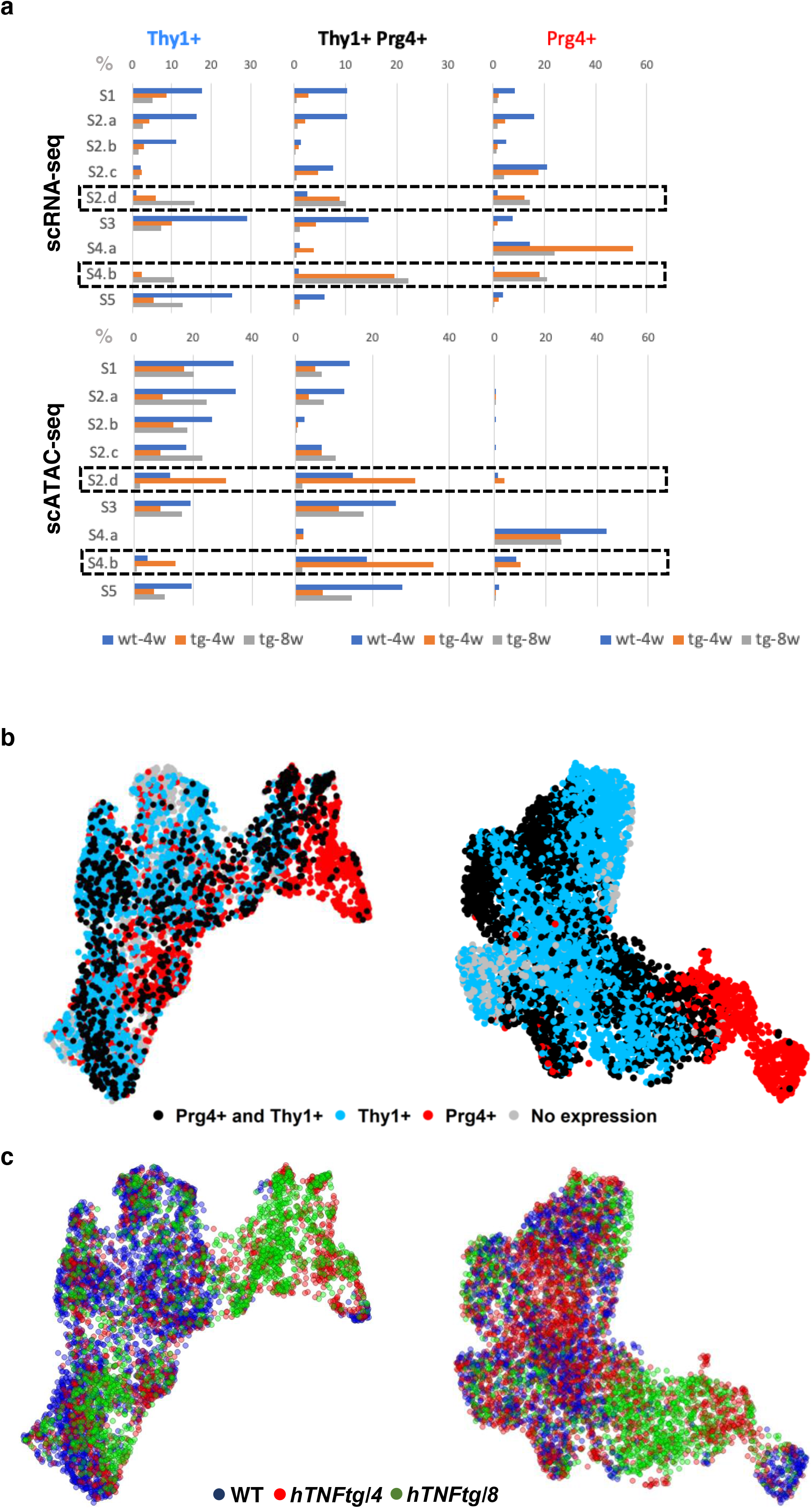
Prg4 and Thy1 expression in hTNFtg and WT SFs. (a). Barplots showing the percentage of Thy1 positive, Prg4 positive and double positive SF cells in each cluster per sample for scRNA-seq (top) and scATAC-seq (bottom) modalities. (b). UMAP showing SFs distribution for the pooled scRNA-seq (left) and scATAC-seq (right) datasets. Cells are colored for Thy1 positive, Prg4 positive, double positive SF cells, and double negative cells. (c). Similar to (b), but cells are colored by sample of origin.

**Supplementary Figure 5.**
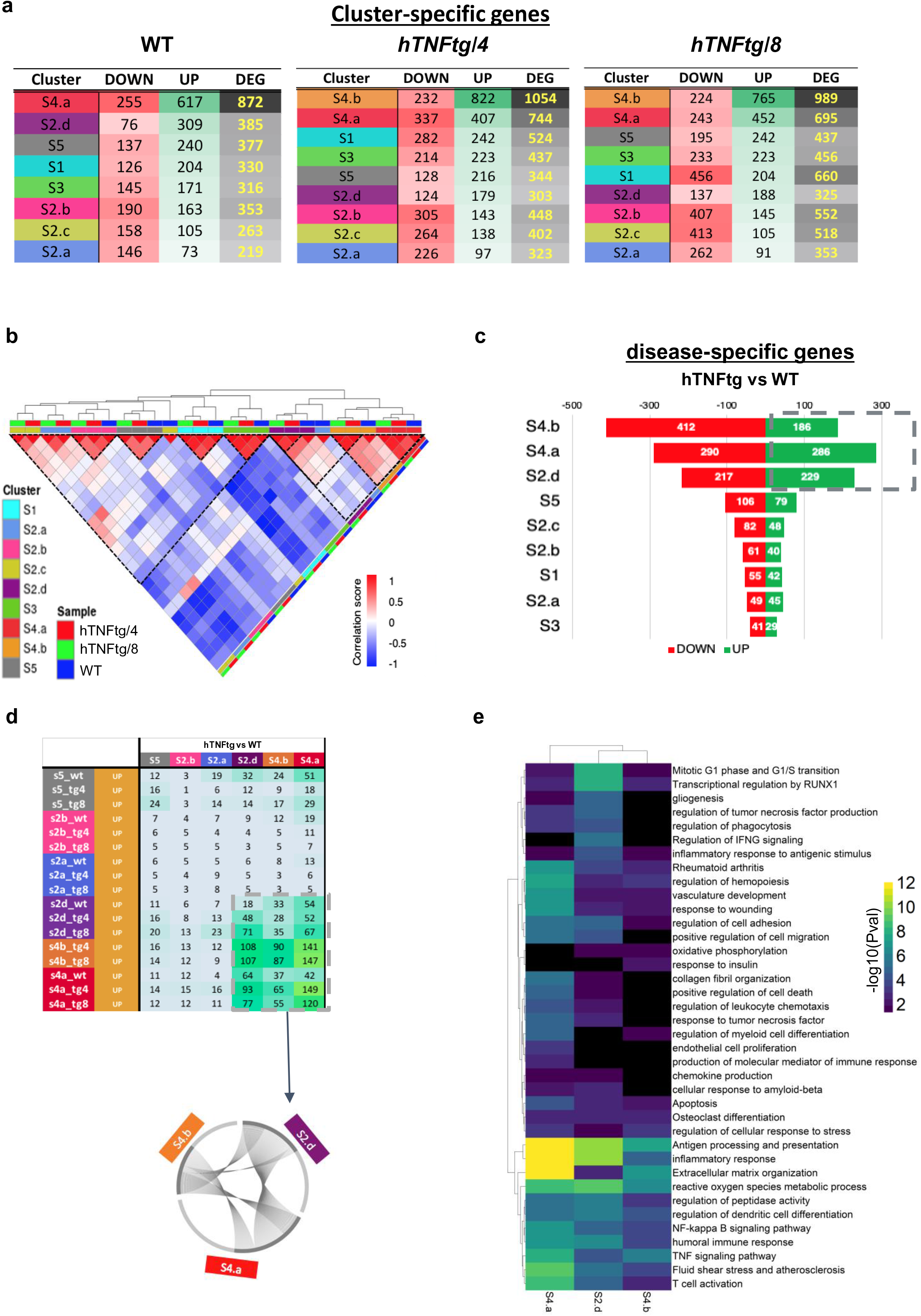
Differential Expression Analysis (DEA) of hTNFtg and WT SFs transcriptomes. (a). (Top) Tables summarizing the numbers of up-regulated (UP) and down-regulated (DOWN) genes after DEA between a given cluster and the others (inter-DE) for WT, Tg4 and Tg8 cells. (b). Heatmap of correlation scores between most variable genes (MVGs) for all SF clusters in WT, hTNFtg/ 4w and hTNFtg/ 8. Individual clusters and samples are color-coded. Pathology- and homeostasis-specific highly correlated groups of clusters are highlighted with black dotted lines. (c). Divergent barplot showing the number of up (green) and down (red) -regulated genes after DEA between Tg and wt cells (Intra-DE) of a given cluster. (d). (Top) Table summarizing the number of inter-intra DE genes commonly up-regulated in different cell subsets showing the similarities in genes defining S2.d S4.b and S4.a clusters (doted box) and (Bottom) circos plot visualizing the extent of shared up-regulated genes between these clusters. (e). Functional enrichment analysis of intra-DEGs (hTNFtg vs WT) for the clusters S2.d, S4.b and S4.a. Functional terms in the heatmap are colored according to statistical significance of the enrichment.

**Supplementary Figure 6.**
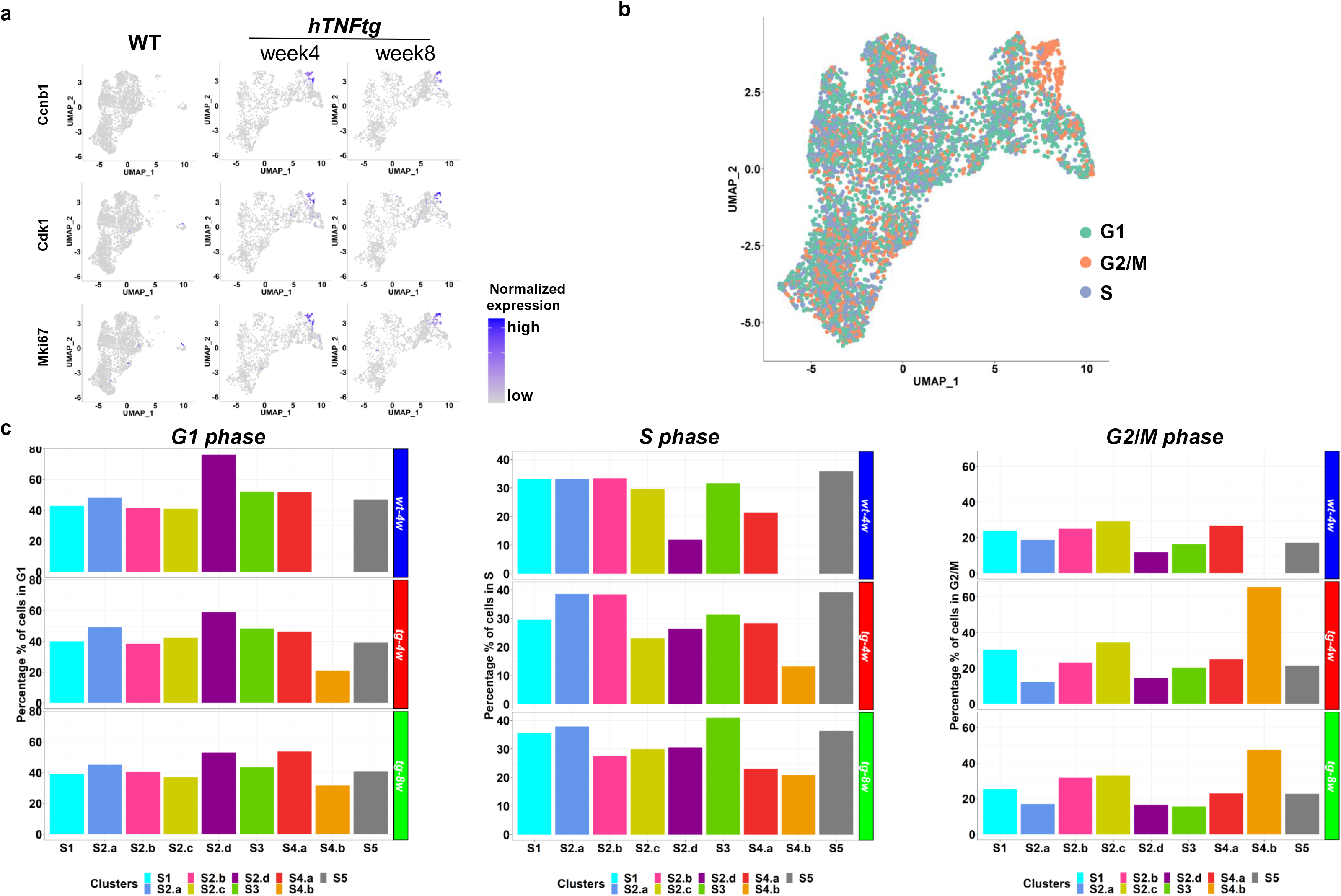
Cell cycle analysis of SFs from hTNFtg and WT mice. (a). Feature plots showing normalized expression values of genes associated with cell cycle proliferation. (b). UMAP projection depicting the phase of the cell cycle for each cell in the pooled dataset (see Methods). (c). Barplots showing the percentages of cells in each cluster (per sample) belonging to phase G1(left), S(center) and G2/M(right).

**Supplementary Figure 7.**
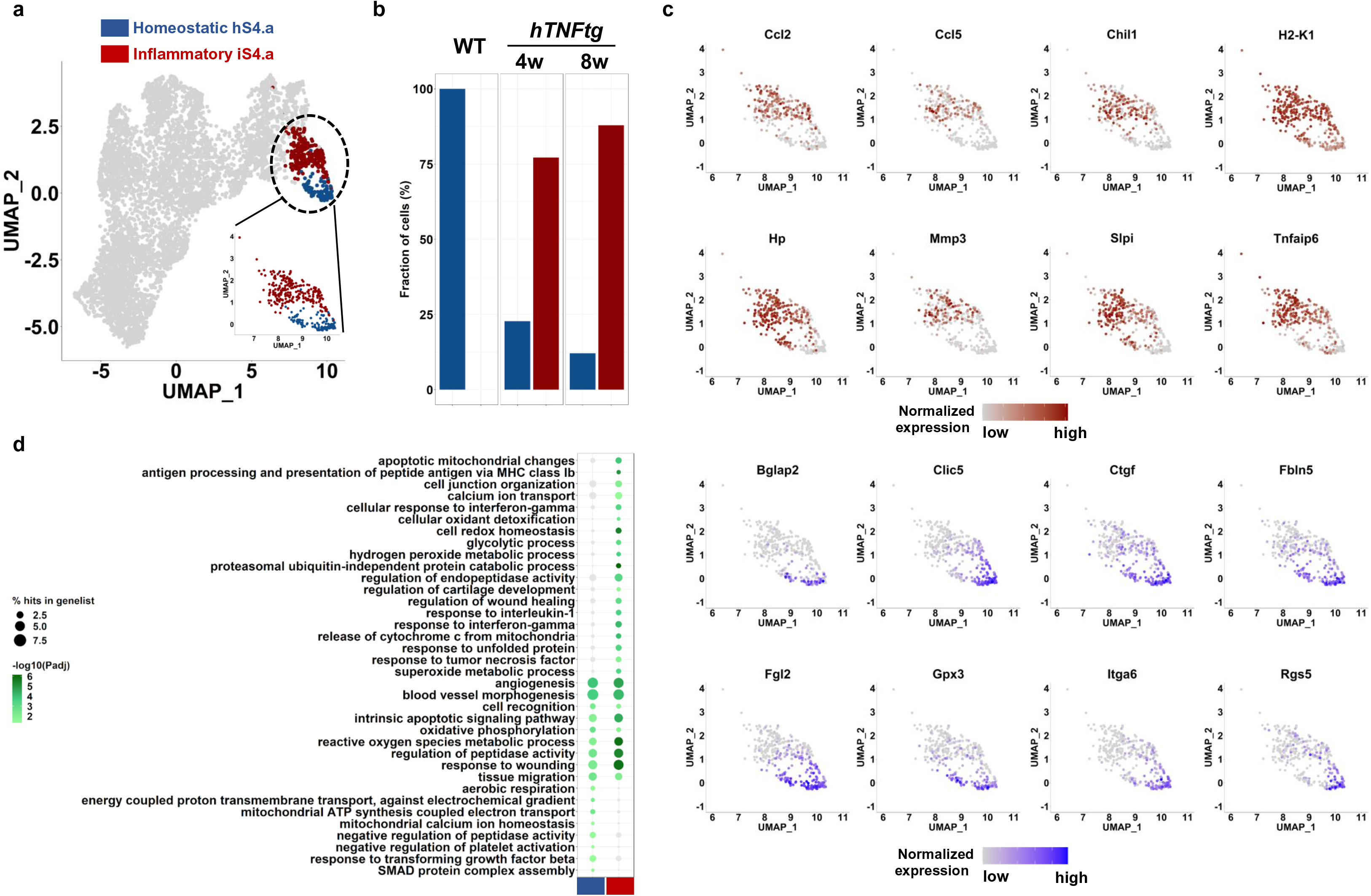
Sub-clustering analysis of lining population revealed two distinct sub-clusters. (a). UMAP representation of the identified lining sub-clusters. Cells belonging to the homeostatic group are colored in blue, while cells belonging to the inflammatory group are colored in red. UMAP plots of the re-clustering is also shown as an inset. (b). Barplot showing the relative abundances of cells in each sub-cluster in healthy condition and during disease progression. (c). UMAP Feature plots of selected genes showing differential patterns of expression between the two sub-clusters. (d). Dotplot of shared and specific enriched GO biological processes in the two sub-clusters.

**Supplementary Figure 8.**
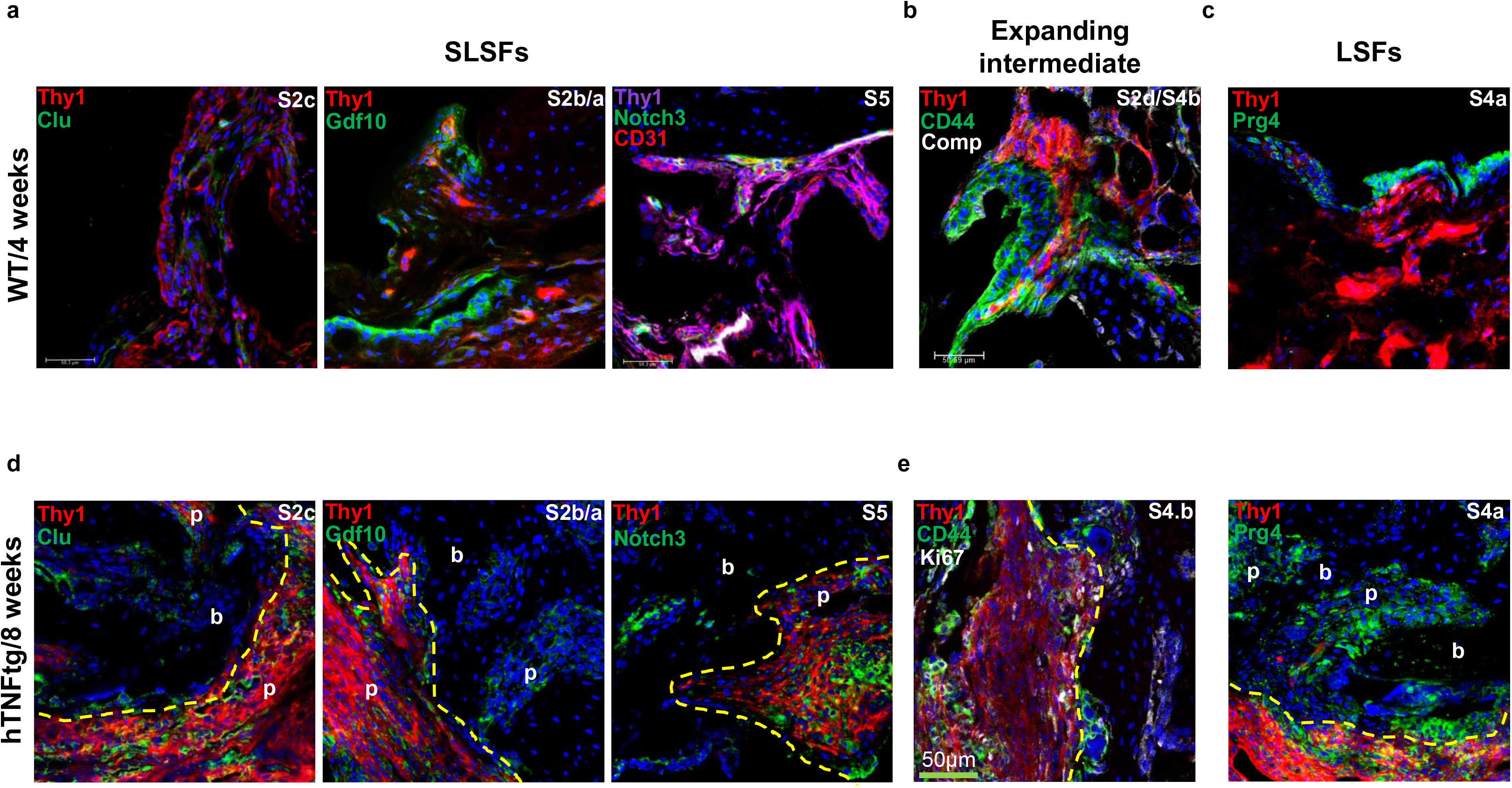
Spatial distribution of representative sc SFs clusters markers in the invasive pannus formed at hTNFtg ankle joints at an established arthritis state. Representative confocal section (n=3 mice per genotype) depicting expression of marker genes identified upon SF clustering. Dashed yellow line in all the hTNFtg images indicates, sub-lining localization at the level of the intersection between pannus (p) and bone (b), dashed lines in every picture and highlights the destructive pannus that has invaded in the talus bone of the ankle joint. Representative confocal images of ankle joint sections at the talus level of 4-week-old wt mice, top panels and 8-week-old hTNFtg mice (n= 3 mice per genotype). The marker genes and the associated subpopulations are indicated on each panel. (a)., (d). S2c marker Clu shows a well distributed expression withing the sublining compartment, defined by Thy1 expression. S2a and b marker Gdf10 in green, shows expression mainly in the pro-inflammatory pannus surrounding the joint. S5 marker Notch3 displays a perivascular localization in the healthy synovial membrane, and an expansion in the arthritic ankle joint, co-localized withing the Thy1+ sublining. (b)., (e). Cd44 in green and Comp in white co-expressed in the restricted S2d subpopulation in the healthy joint, however, both expanding arthritic subpopulation S2d and S4b indicated by co-staining of Cd44 with Ki67, are localized at the edges of the pannus both inside and outside of the bone. (c)., (f). Prg4 in green, marker gene for the S4a lining subpopulation displays exclusive expression in the outermost SF layer of the healthy joint and in the hTNFtg joint, its expression is expanded and is mainly detected in the invasive pannus in the bone but also within the interface pannus bone zone. All Scale bars, 50μm., b: bone, p: pannus.

**Supplementary figure 9.**
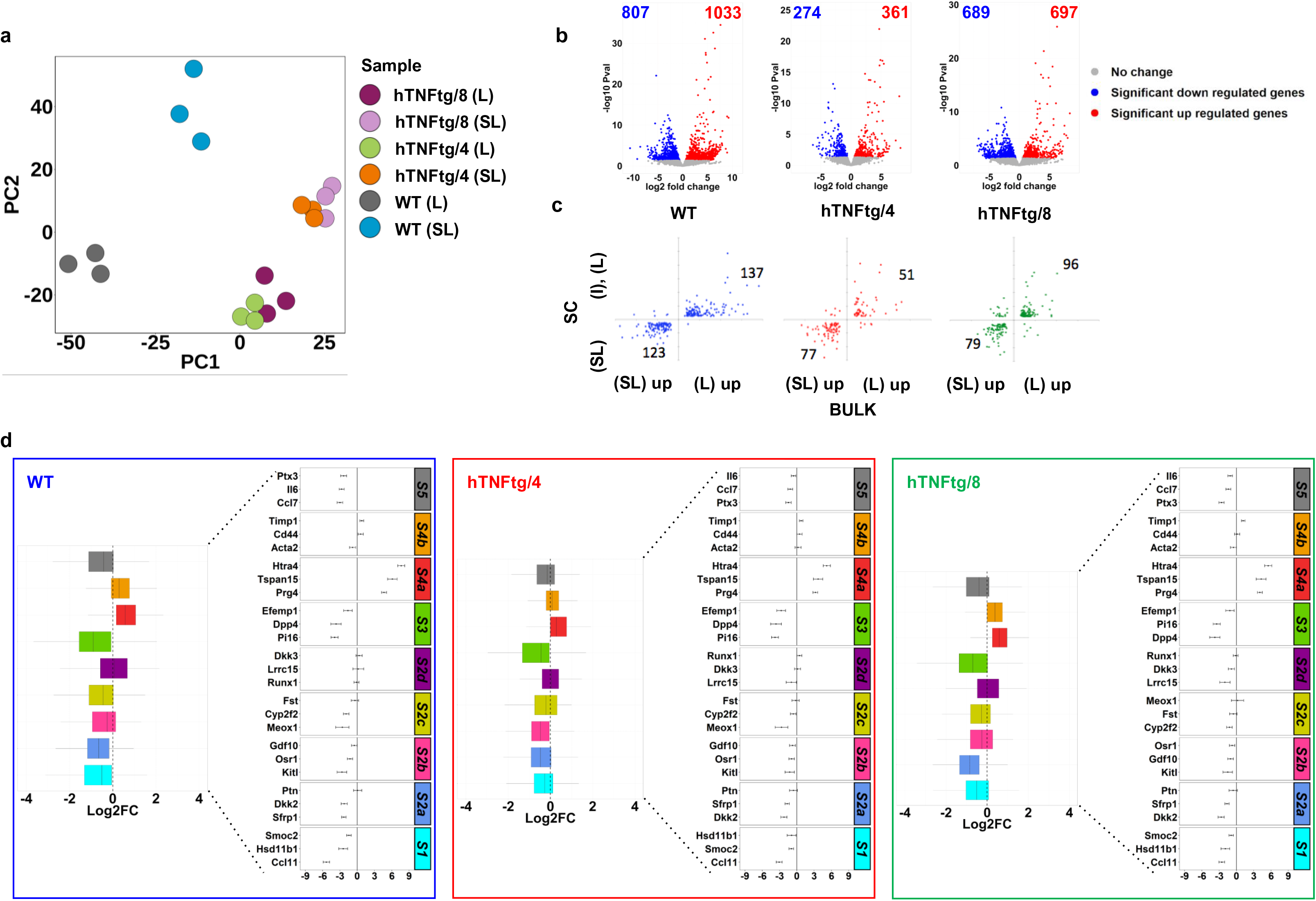
Bulk RNA-seq analysis of sorted SFs from ankle joints of WT and hTNFtg mice (1/2). (a). PCA plot of bulk RNA-sequencing samples. Note how samples are grouped horizontally as wild type (left) and transgenic (right) and vertically as sub-lining (top) and lining (down) (b). Volcano plots showing the number and characteristics of up and down regulated genes for the comparisons WT L vs WT SL, Tg4 L vs Tg4 SL and Tg8 L vs Tg8 SL (c). Scatter plot showing the extent of the overlap (and the correlation in FCs) for lining, intermediate and sublining specific DEGs between scRNA-seq and bulk RNA-seq. Note the lower numbers in hTNFtg samples. (d). On the left side: box plots showing Log2FC values(L vs SL comparisons) in each sample of bulk RNA-seq data for the marker genes of each cluster in the scRNA-seq dataset. On the right-side, error-bar plots showing the standard error of Log2FC values for 3 selected marker genes per cluster. Results are shown per sample as indicated.

**Supplementary figure 10.**
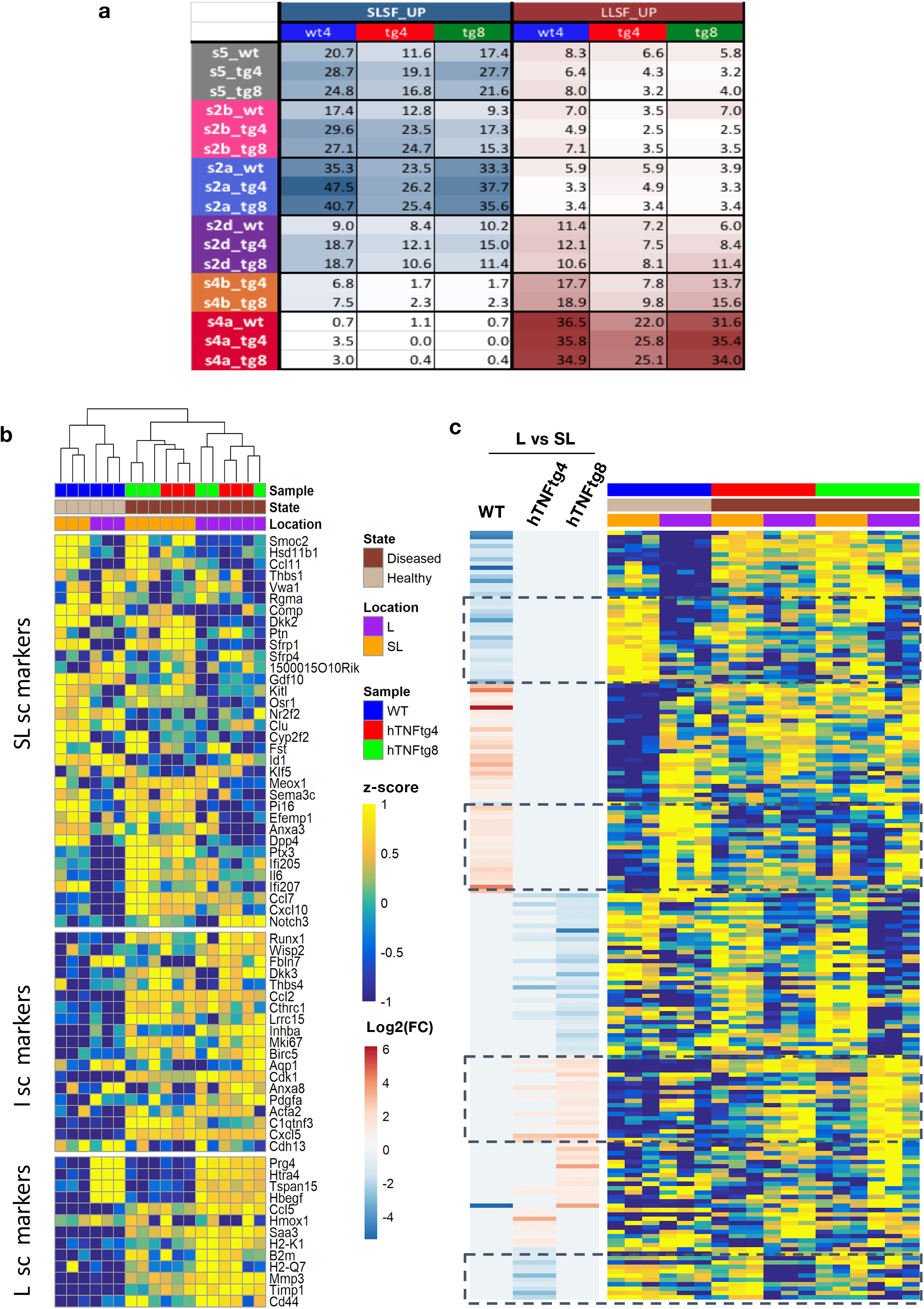
Bulk RNA-seq analysis of sorted SFs from ankle joints of WT and hTNFtg mice (2/2) (a). Proportions (%) of the marker genes (found by scRNA-seq for a specific cluster), which are also up regulated in bulk RNA-seq for lining (shades of red) and sublining cells (shades of blue). (b). Heatmap of scaled normalized counts obtained with bulk RNA-seq assay for the sc marker genes described in figure 2C (for sub-lining (SL), Intermediate (I), and Lining (L)). (c). Heatmap showing the differences across samples (compare WT with hTNFtg) in significance of FC between lining (L)and sublining (SL) cells detected in bulk RNA-seq. The boxes highlight candidate genes that can be used for RT-PCR to test disease status.

**Supplementary figure 11.**
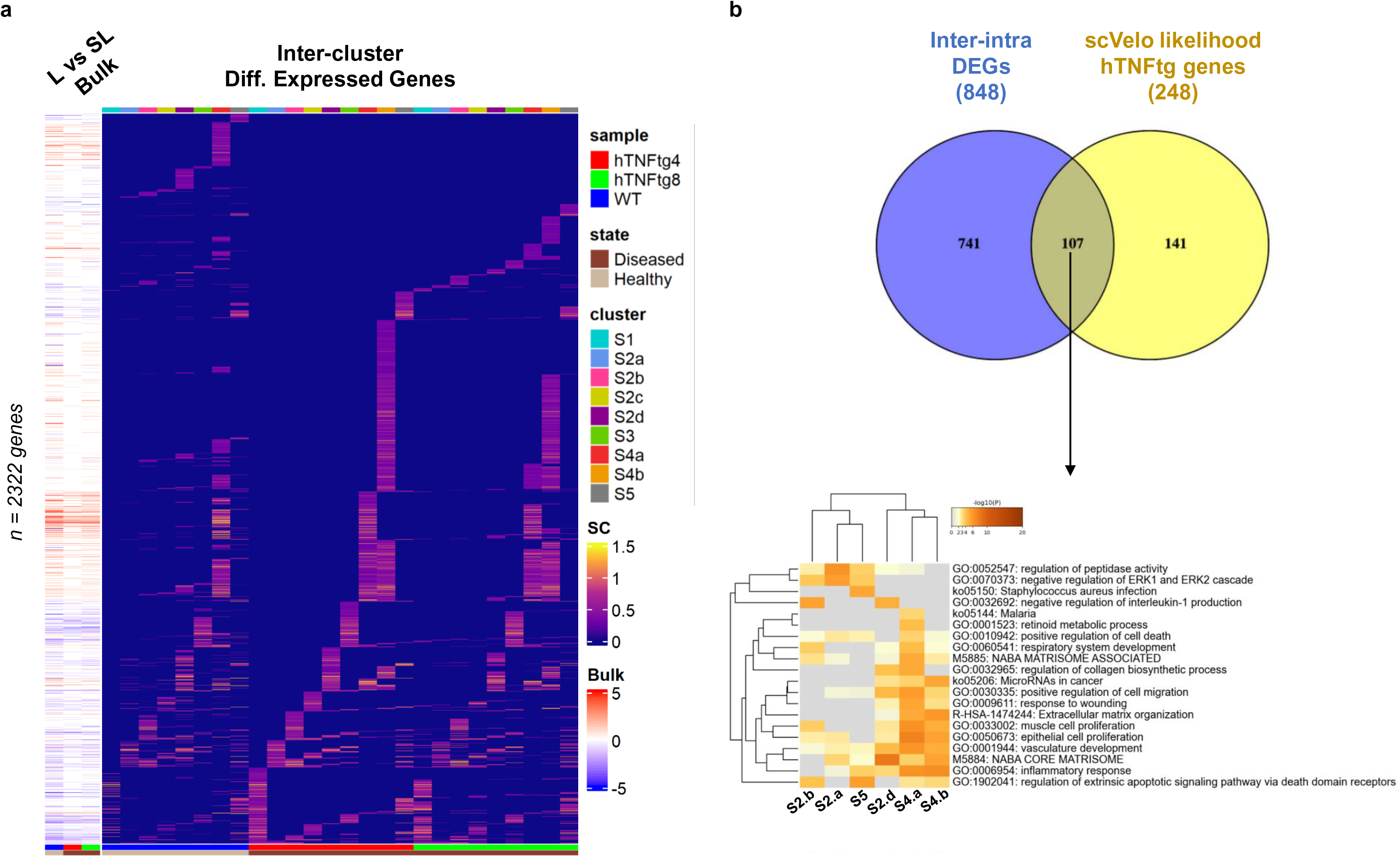
Complementary DE analyses. (a). Right: Heatmap depicting the inter-cluster FC values for the 2322 genes displaying cluster specificity (marker gene of at least one of the represented cluster). Left: Heatmap depicting the bulk RNA-seq FC values of those genes (see Methods). Samples are individually shown. Note the overrepresentation of genes expressed in the lining cells according to bulk RNA-seq in sc S4.a cluster, and the enrichment of genes expressed in the sublining cells according to bulk RNA-seq in homeostatic clusters S5, S2.a and S2.b. In contrast cluster S2.d and S4.b are specified by genes not DE between join compartments according to bulk RNA-seq (see Supplementary figure 10a for quantification). (b). Top: Venn diagram showing the extent of the overlap (107 genes) between inter-intra DEGs and scVelo likelihood hTNFtg genes for the clusters of the trajectory path described in Figure 5. Bottom: Heatmap showing clusters obtained after Functional enrichment analysis. Note the co-clustering of pathogenic (S2.d, S4.b, S4.a) cells functions.

**Supplementary figure 12.**
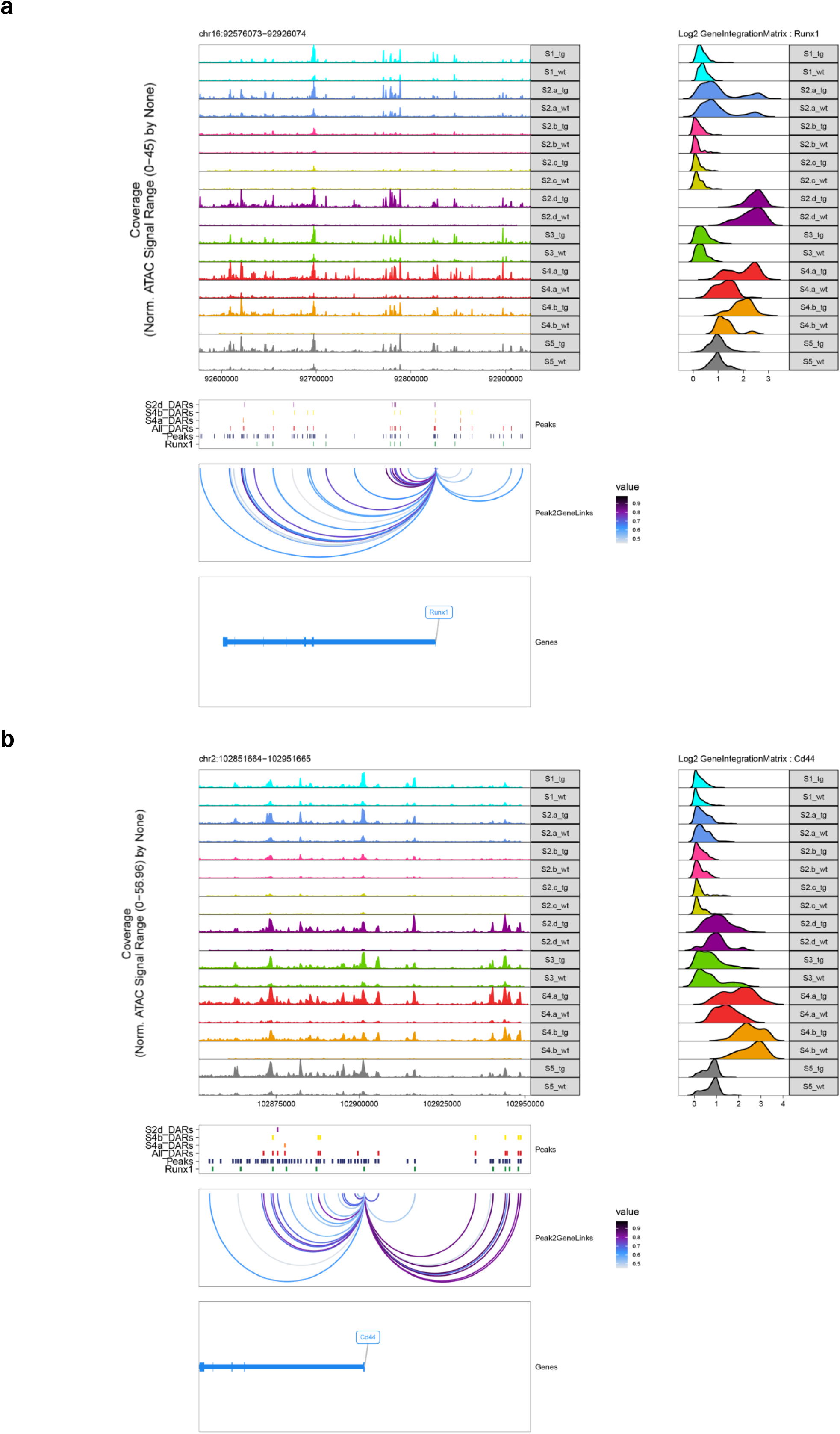
Runx1 motif-enriched regulatory elements control the expression of Runx1 and Cd44 genes during the arthritic disease state. (a). Genome track snapshot of the extended Runx1 gene locus (chr16, 92,576,073–92,926,074). Single-cell gene expression (Log2 GeneIntegrationMatrix: Runx1) across SF subpopulations and disease states (wt, hTNFg) is shown to the right. Merged peaks across SF subpopulations (Peaks), and differentially accessible peaks between SF subpopulations (All_DARs) are shown below. Differential accessible peaks with significantly increased scATAC-seq signal in S2d hTNFg (S2d_DARs), S4b hTNFg (S4b_DARs), and S4a hTNFg (S4a_DARs) cells are also reported accordingly. Inferred peak-to-gene linkages for intragenic and distal intergenic regulatory elements are shown below (Peak2GeneLinks). Color scale signifies the level of correlation between peak scATAC-seq accessibility and integrated gene scRNA-seq expression (value). (b). same as in a for the extended Cd44 gene promoter region (chr2, 102,851,664– 102,951,665).

**Supplementary Figure 13.**
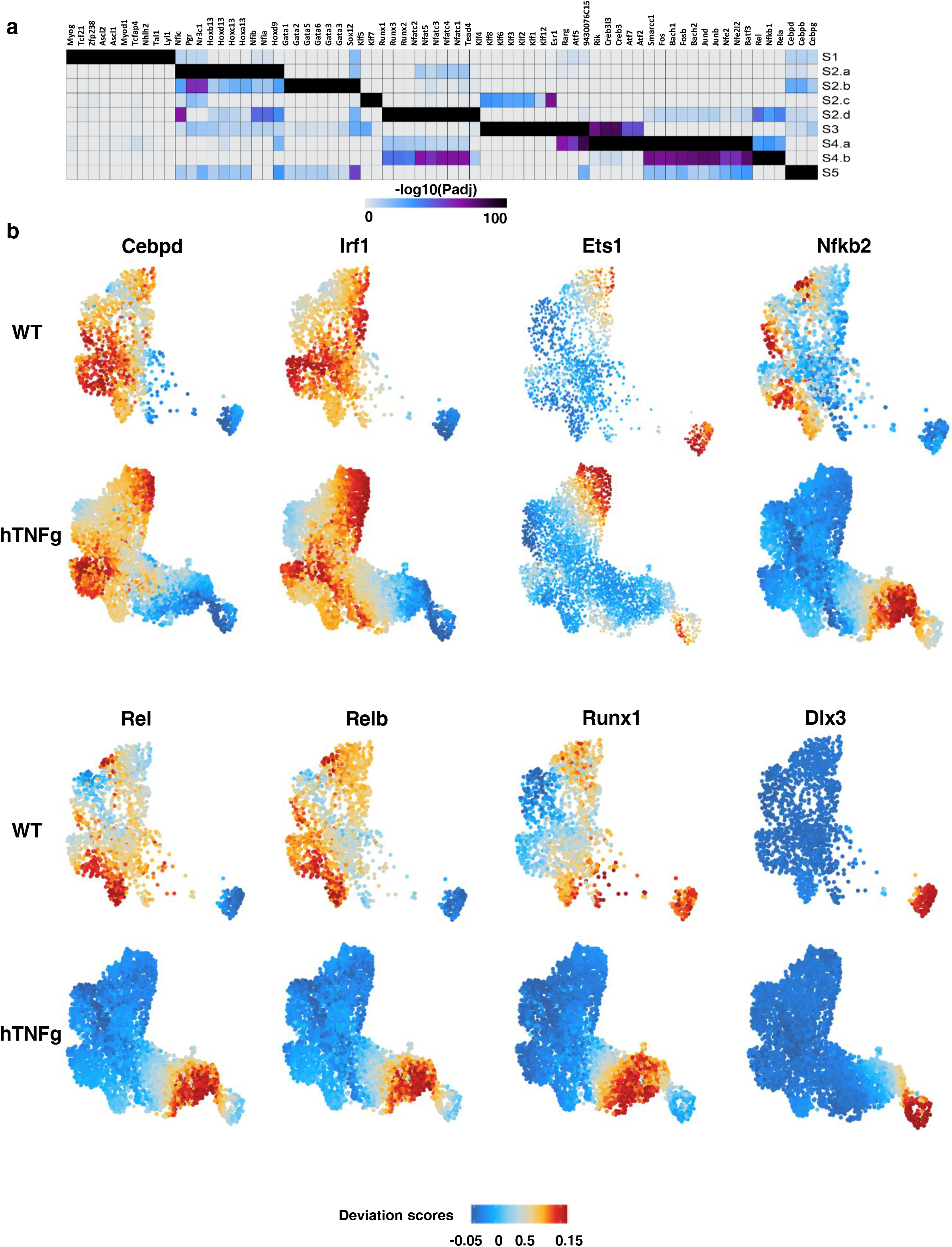
Motif enrichment analysis in scATAC-seq cluster-specific accessible regions define distinct types of regulatory programs across SF subpopulations. (a). Heatmap showing the motif enrichment P-adjusted values of each SF subpopulation. Motif enrichment analysis was performed within the SF marker peaks depicted in figure 1D (lower panel). Color signifies the magnitude of the enrichment (−log10 (P adjusted value), hypergeometric test). Columns are order by using binary sorting. (b). Feature plots of selected TF motifs with regulatory activity in the homeostatic, intermediate and lining SF subpopulations. Color signifies the motif deviation scores (see Figure 4 and Methods).

**Supplementary Figure 14.**
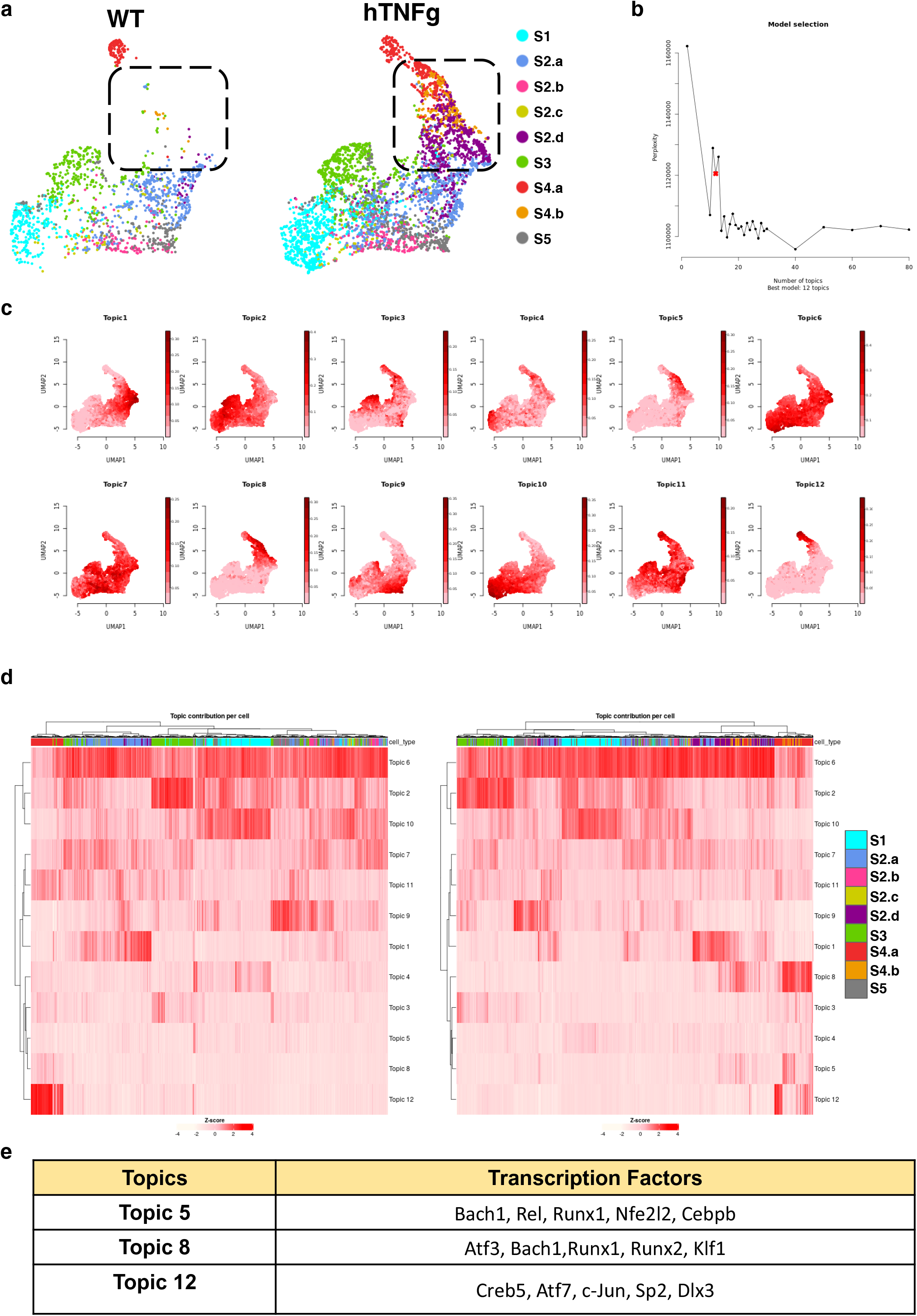
Cis-regulatory modeling (CisTopic) on scATAC-seq recapitulates the identification of putative arthritic regulatory programs. (a). UMAP representation of SFs across the two disease states using cell-topic probabilities. (Wt and hTNFtg as indicated). Cells are colored by cluster identities and the dotted line/marked area highlights the dynamic changes of the intermediate and lining subpopulations during disease progression. (b). CisTopic modelling (see Methods) of cis-regulatory topics using Latent Dirichlet Allocation. An optimal model of 12 topics (arrow) was selected based on log likelihood. (c). UMAP Feature Plots of per-cell topic probabilities for each modeled topic on the aggregated scATAC-seq dataset. (d). Heatmaps showing the per-cell topic z-scores (columns) for each modeled topic (rows), for each disease state (left panel: WT, right panel: hTNFtg). (e). Selection of representative motif enrichment results found on the most contributing regions of topic 5, topic 8 and topic 12.

**Supplementary figure 15.**
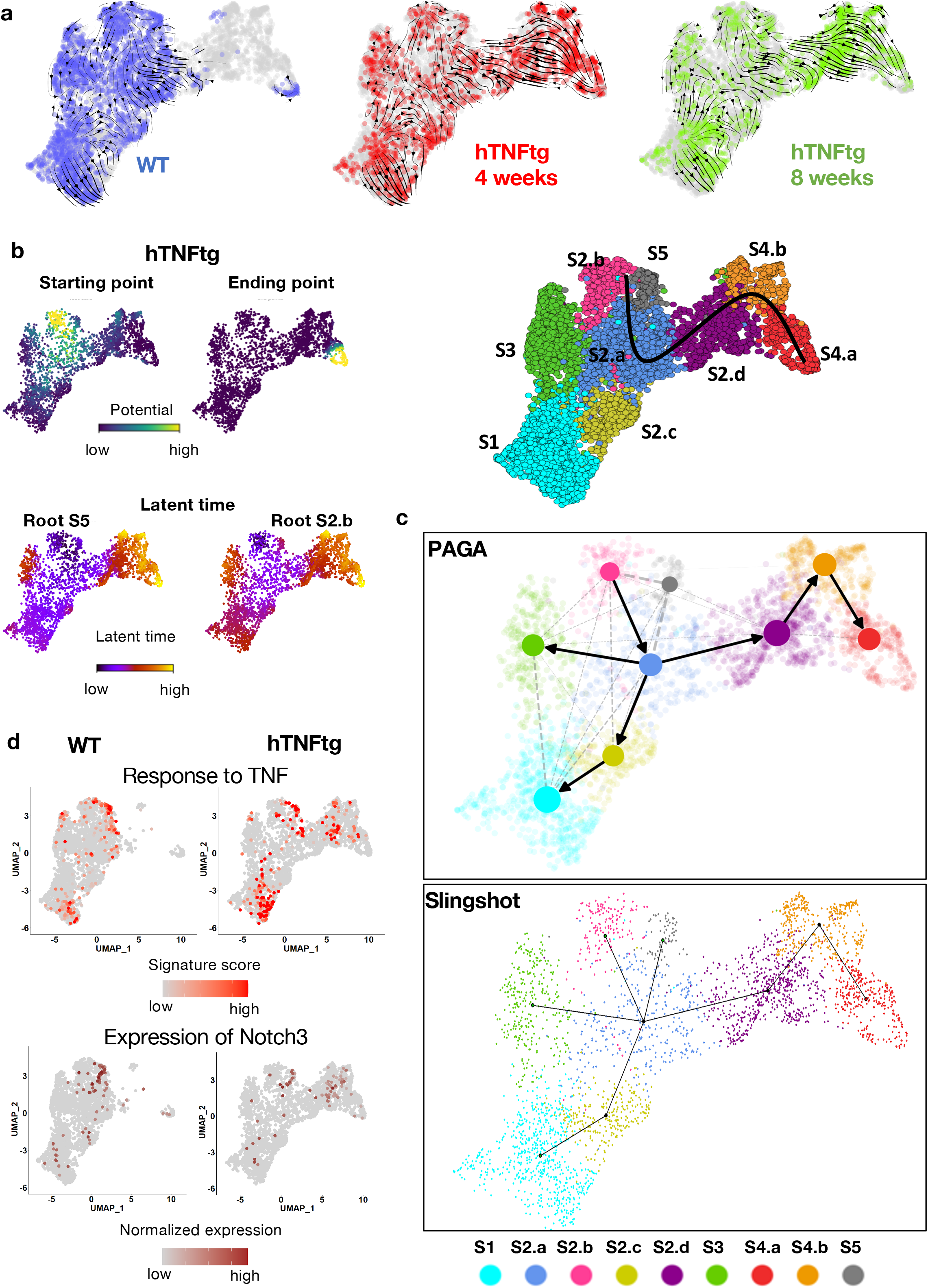
Ordering of the cells across the identified trajectory using latent time values and alternative methodologies. (a). UMAP plots displaying RNA velocity results detailing cell transitions and dynamic relations between SF clusters in the WT and the hTNFtg datasets (week4 and week8, Tg4 and Tg8 respectively). (b). left side: UMAP plost of hTNFtg sample, cells are colored by their potential of belonging to the initial(left) or final state(right) of the identified trajectory (upper panel) and UMAPs of hTNFtg samples, cells are colored by latent time value (lower panel). In the latent time panels, on the left, the calculations were performed by considering the root cell from cluster S5, while on the right panel cells from cluster S2.b were considered root. (See Methods). Right Side: UMAP plot highlighting the selected trajectory trend across S2.b/S5, S2.a, S2d, S4.b and S4.a). (c). Partitions identified by PAGA algorithm and minimum spanning tree produced by Slingshot propose a similar global structure of the mouse data, supporting the existence of a trajectory backbone which includes clusters S2.a, S2.d, S4.b, S4.a. (d). UMAP projections of WT (left) and hTNFtg (right) samples, where cells are colored based on signature scores for the GO term “Response to TNF” (upper panel) or on the normalized expression of Notch3 (lower panel).

**Supplementary Figure 16.**
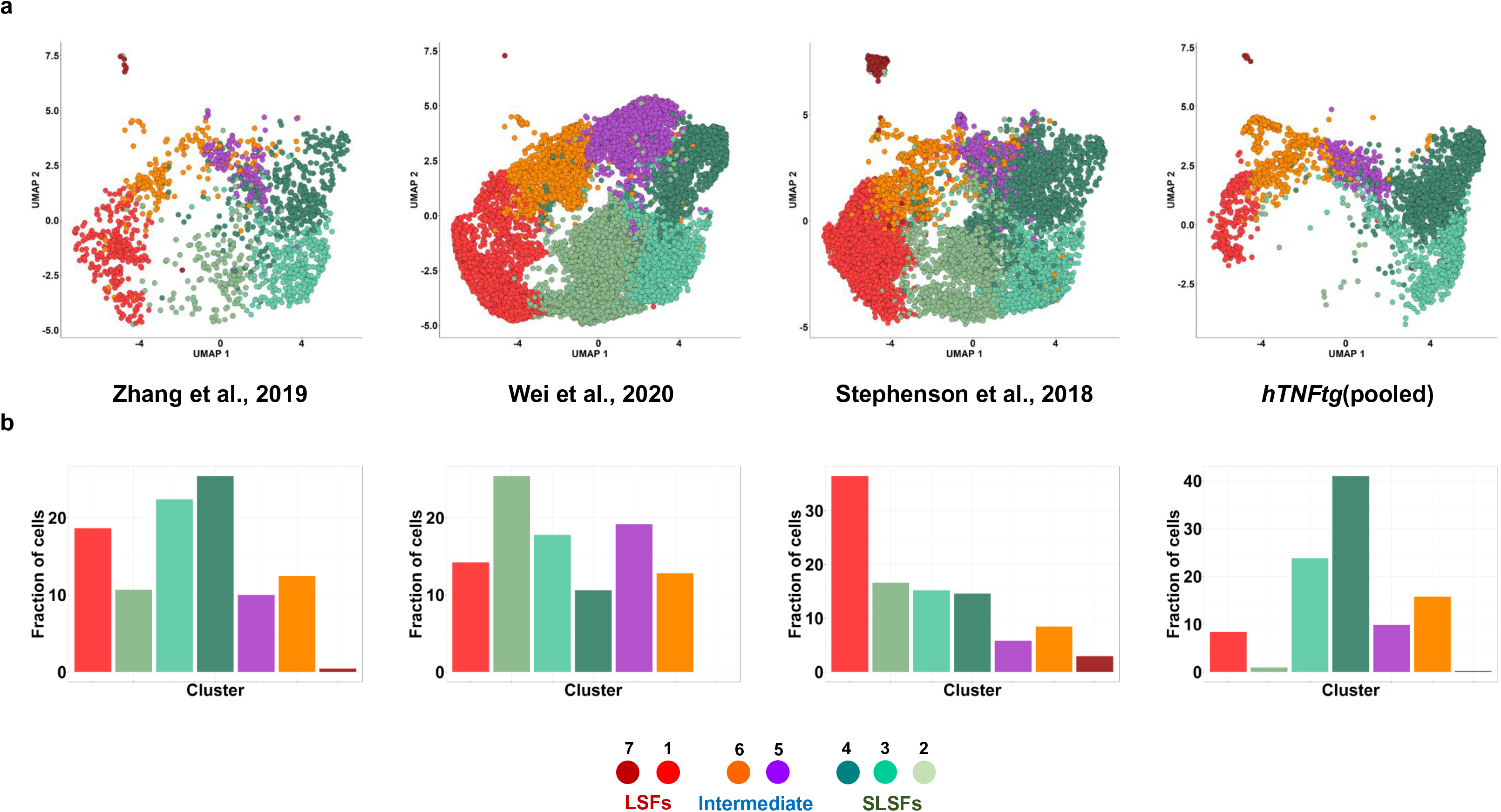
Human/Mouse integrative analysis employing the available scRNA datasets from SFs of RA patients and the hTNFtg mice. (a). UMAP projections showing the distribution of cells in the integrated aligned H-M clusters for each of the datasets used in human mouse integration analysis, as indicated. (b). Barplots showing the relative abundances of cells belonging to each of the integrated clusters in the datasets described in (a).

**Supplementary Figure 17.**
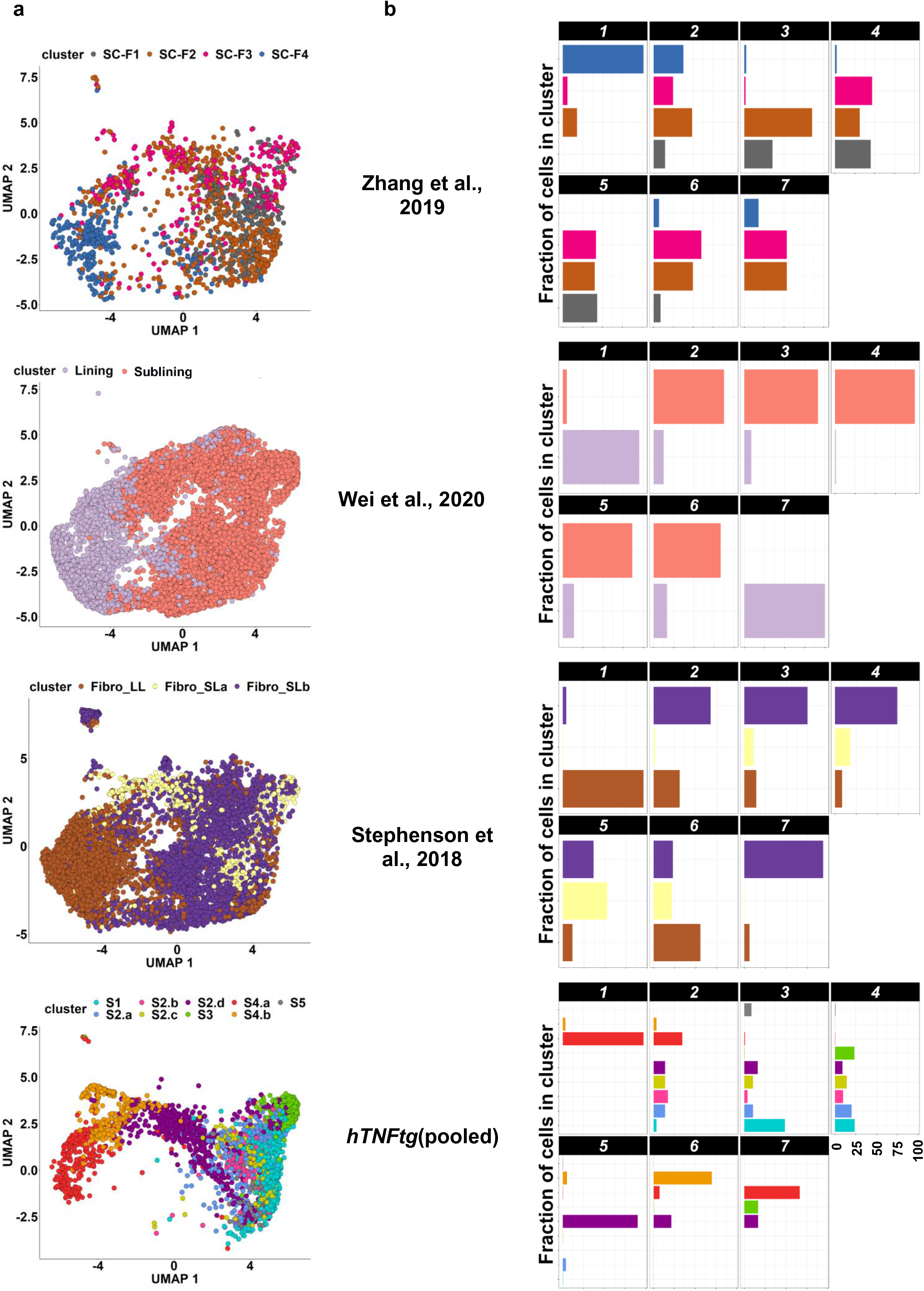
Distribution of human and mouse SF cells as previously annotated upon the integrative analysis. (a). Cells from the 4 datasets used in integration analysis are plotted in the common integrated UMAP space and are colored by their original annotation. (b). Barplots showing the distribution of cells (using the original annotation) to the integrated clusters for the datasets on the left.

**Supplementary Figure 18.**
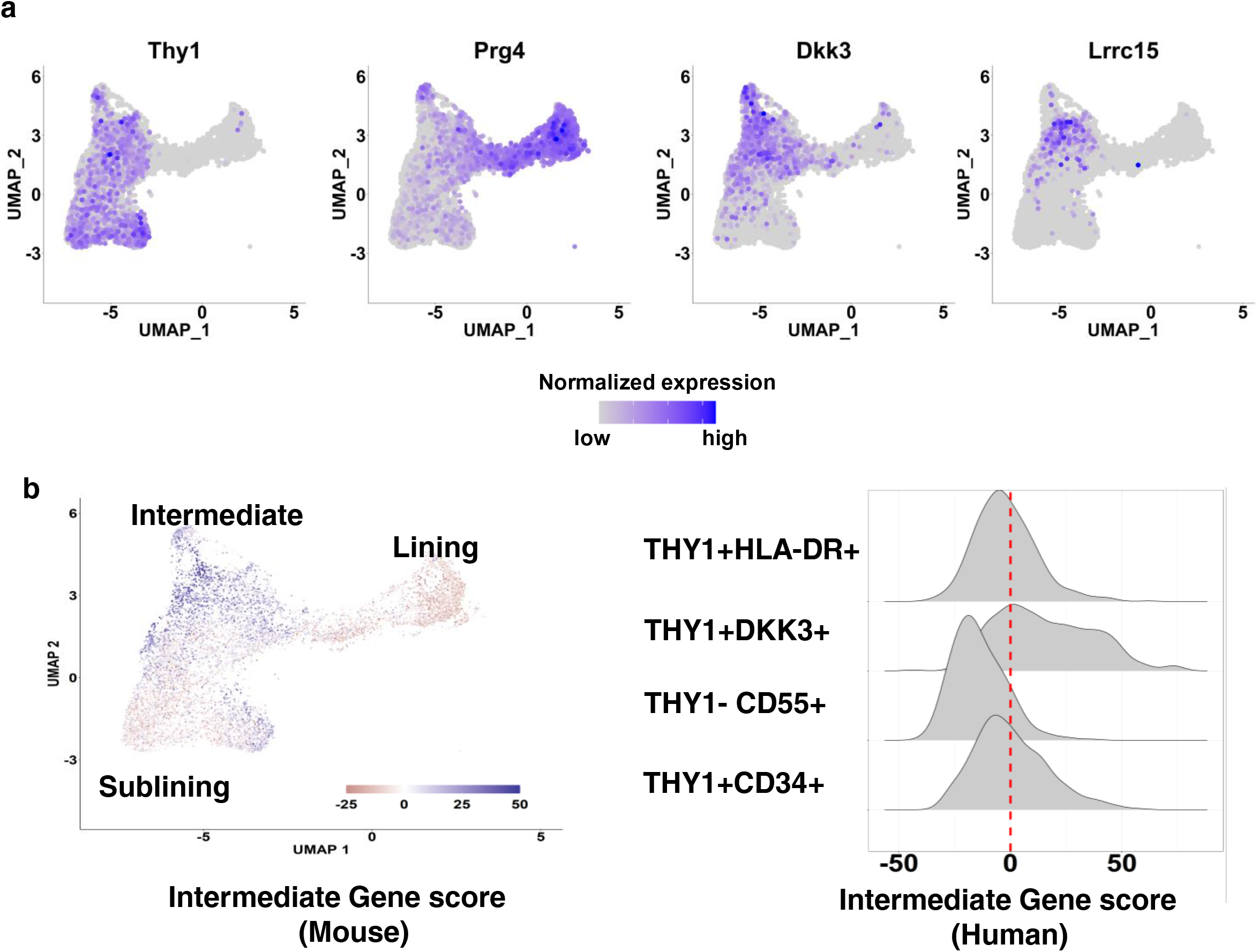
Dkk3 and Lrrc15 expression in the intermediate transcriptional state of SFs in murine and human arthritis. (a). Feature plots of mouse synovial fibroblasts from Wei et al., 2020. Cells are colored by normalized expression of genes Thy1, Prg4, Lrrc15 and Dkk3. (b). UMAP projection(left) of mouse synovial fibroblasts shown in (a). Cells are colored by signature score. The signature score is calculated as the sum of scaled normalized expression values for the 71 intermediate genes described in Wei et al., 2020. Density plot(right) of the same signature scores for human RA synovial fibroblasts from Zhang et al., 2019.

